# Semaphorin7A and PD-L1 cooperatively drive immunosuppression during mammary involution and breast cancer

**DOI:** 10.1101/2025.01.03.631229

**Authors:** Alan M Elder, Heather R Fairchild, Kelsey T Kines, Lauren M Cozzens, Alexandria R Becks, Traci R Lyons

**Affiliations:** Division of Medical Oncology, Department of Medicine, School of Medicine, University of Colorado Anschutz Medical Campus, Aurora, CO, USA; Cancer Biology Graduate Training Program, University of Colorado Anschutz Medical Campus, Aurora, CO, USA; Medical Scientist Training Program, University of Colorado Anschutz Medical Campus, Aurora, CO, USA; Young Women’s Breast Cancer Translational Program, University of Colorado Anschutz Medical Campus, Aurora, CO, USA; University of Colorado Cancer Center, Aurora, CO, USA

## Abstract

Postpartum mammary gland remodeling after a pregnancy/lactation cycle is characterized by mechanisms of immunosuppression. Here we show that SEMA7A promotes PD-L1 expression in immune cells of the mammary tissue during involution. These same phenotypes are mimicked in the microenvironment of SEMA7A-expressing tumors, which partially respond to αPD-1/αPD-L1 treatments in vivo. However, cells that remain after treatment are enriched for SEMA7A expression. Therefore, we tested a novel monoclonal antibody that directly targets SEMA7A-expressing tumors, in part, by reducing SEMA7A-mediated upregulation of PD-L1. In vivo, the SEMA7A monoclonal antibody also reduces tumor growth or promotes complete regression of mouse mammary tumors, reduces the immunosuppressive phenotypes in the tumor microenvironment and restores cytotoxic T cells suggesting that SEMA7A may be a candidate for a novel immune-based therapy for breast cancer patients.

## Introduction

The protective effect of pregnancy and breast feeding against development of breast cancer has been described, particularly in women that have their first child at a younger age^1–8^. However, there exists a transient increased risk for developing breast cancer that is associated with childbirth at any age and the risk window for developing breast cancer is highest in women that have their first child after 35 years of age^4,5,9^, a practice that is trending upward globally. Postpartum breast cancers (PPBC) have been defined as breast cancers diagnosed within a decade of childbirth based on the poor prognosis observed in these patients^10–17^. Specifically, PPBCs are enriched for involved lymph nodes, lymphovascular invasion, luminal B subtype, and recur more frequently than breast cancers in nulliparous women^10,11,18–21^. This devastating diagnosis affects thousands of young women annually.

Mouse models have revealed that the process of postpartum mammary gland involution can promote both tumor cell growth and metastasis in xenograft and isograft models. In rodents, the process of postpartum mammary involution is characterized by death of 80% of the secretory mammary epithelial cells that arose during pregnancy to support milk secretion during lactation; the associated remodeling of the gland to a pre-pregnant-like state has multiple similarities with wound healing^22–27^. During the first phase of involution, studies in mice, cattle, and sheep have shown that there is an influx of neutrophils, macrophages, and eosinophils that help prevent infections and promote resolution of mastitis^28–35^. During the second phase of involution, macrophages and lymphocytes are increased and have roles in suppressing inflammatory responses in the involuting mammary gland to allow for active tissue remodeling^22,23,36–39^. In additional published studies, lymphatic vessels are increased, M2 polarized macrophages are required, a subset of M2-like macrophages termed podoplanin (PDPN) expressing macrophages, or PoEMs^40^, are present, and increases in T-regulatory cells are observed^20,22,38,41^. Additionally, lymphatic endothelial cells (LECs) and macrophages that express programmed death ligand-1 (PD-L1), and programmed death-1 (PD-1) positive T cells are more frequent in the mammary gland during involution^42^. PD-1 binding of PD-L1 is a known signaling mechanism for T cell exhaustion in the tumor microenvironment (TME) and these same phenotypes are observed in pre-clinical models of PPBC where it was shown that tumor growth after implantation during postpartum involution can be abrogated by targeting PD-L1, which also increases cytotoxic CD8+ T cell activation^42^. Finally, multiple aspects of immunosuppression are employed throughout involution in the mammary tissue and studies have identified a similarly “imprinted” immune milieu in mouse models of PPBC, with some validation in human tissues^17,18,37,38,43–45^, suggesting that these immune suppressive phenotypes may contribute to the poor prognosis observed in patients.

Tumors implanted into the mammary gland during involution display increased tumor growth compared to tumors in nulliparous mammary glands and previous studies have shown that semaphorin 7a (SEMA7A) is necessary and sufficient for this growth promotion^20,46–48^. SEMA7A is a signaling molecule involved in multiple aspects of development and is expressed on the mammary epithelium during involution where it promotes cell survival, in part, via alterations to AKT mediated pro-survival signaling^25–27,46,47,49,50^. In addition, SEMA7A expression is higher in tumors that form in postpartum mice, is more highly expressed in PPBCs compared to tumors in age/stage matched nulliparous women^20,21^ and can predict for recurrence in PPBCs. Pleiotropic roles for SEMA7A in multiple aspects of tumorigenesis have been identified including increased cell viability, migration, invasion, epithelial to mesenchymal transition (EMT), lymphangiogenesis, macrophage infiltration, macrophage lymphatic interactions, fibroblast deposition of collagen, and metastasis in mouse models ^20,46–48,50–53^. Additionally, SEMA7A expression drives decreased distant metastasis free survival in patients including those whose tumors are estrogen receptor positive (ER+) and estrogen receptor negative (ER-) breast cancers^47^. Collectively, these data suggest that SEMA7A drives tumor progression in both PPBCs and breast cancer in general. As such, identification of therapeutic strategies for targeting SEMA7A+ tumors should be explored for improving patient survival. In this manuscript, we explore the relationship between SEMA7A and targets of immunotherapy, namely PD-1 and PD-L1, which could be exploited with currently available, FDA approved, immunotherapies and/or with direct targeting of SEMA7A.

Anti-PD-1 and anti-PD-L1 (αPD-1/αPD-L1) based therapies are some of the best advances in immuno-oncology; these antibody-based therapeutics have demonstrated great success clinically for multiple tumor types including melanoma, head and neck, and lung cancers^54–63^. One αPD-1/αPD-L1 therapy that targets PD-1, pembrolizumab (Keytruda), has shown efficacy in patients with advanced triple negative breast cancer (TNBC)^64–66^; however, αPD-1/αPD-L1 therapies have been less successful in breast cancers compared to other cancer types. Thus, identification of markers that could predict for PD-1/PD-L1 signaling in breast cancer could also predict for breast cancers that may respond better to αPD-1/αPD-L1 therapies. In this study, we show that tumor cells implanted into SEMA7A knockout (KO) mice (*Sema7a^tm1Alk^*) during involution do not exhibit the same accelerated tumor growth as those implanted into wild-type and reveal SEMA7A-dependent expression of PD-L1 on immune and stromal cell populations in the mammary gland during involution. We also examine the relationship between SEMA7A and PD-L1 in tumor and stromal cells and show that αPD-1/αPD-L1 treatment can slow the growth of SEMA7A expressing tumors. Finally, we show that a monoclonal antibody that targets SEMA7A can result in tumor regression and reactivation of anti-tumor immunity.

## Results

### SEMA7A-mediated tumor growth and PD-L1 expression during postpartum mammary involution

To test the hypothesis that host-derived SEMA7A is necessary to drive tumor growth during postpartum involution, we implanted E0771 wild-type (WT) mouse mammary tumor cells, which have low endogenous expression of SEMA7A^20^, into intact mouse mammary fat pads of wild-type (WT) C57BL/6 and SEMA7A knockout (KO) (*Sema7a^tm1Alk^*/J^67^) mice at involution day 1. We observed increased tumor growth during involution in the WT mice compared to KO (Fig. 1A). Then, to determine whether a single copy of *Sema7a* was sufficient to restore accelerated tumor growth during involution, we implanted E0771 cells into heterozygous (Het) or KO mice that were nulliparous or one day post wean (day 1 of involution); we observed enhanced tumor growth in both the nulliparous and the postpartum Het Mice in comparison to the KO (Fig. 1B, SFig. 1A). Next, to test the hypothesis that SEMA7A is involved in promotion of the tissue microenvironment observed during postpartum mammary gland involution, we harvested whole mammary extracts from WT and KO mice at involution days 2, 3, 4, 6, and 8 and analyzed immune and non-immune cell populations by flow cytometry. The most consistent changes observed in the KO mice occurred at involution days 3 and 6, where decreased lymphatic endothelial cells (LECs), macrophages, and PoEMs were observed, data which is consistent with previous findings that SEMA7A expression is increased as early as day 3 of involution (SFig. 1B-F)^20,50^. PD-L1 expression was examined on these cell populations by flow cytometry to show that the mammary glands from KO mice had decreased PD-L1+ LECs (Fig. 1C), PD-L1+ total macrophages (Fig. 1D), PD-L1+ M2 macrophages (Fig. 1E) and PD-L1+ PoEMs (Fig. 1F). These studies suggest that expression of SEMA7A in the mammary gland during involution contributes to the changes in the tissue microenvironment.

**Figure 1:**
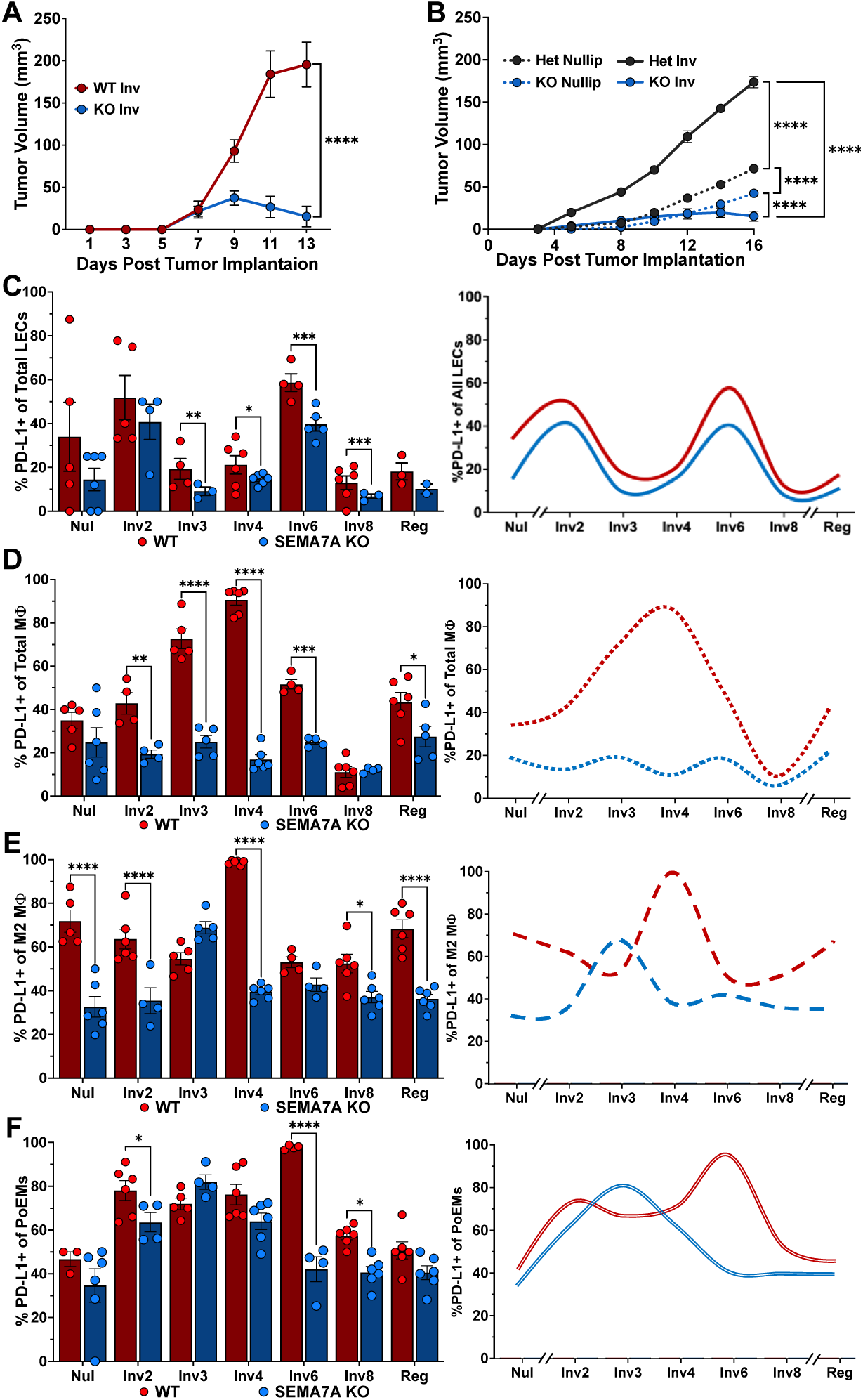
SEMA7A mediates tumor growth and immunosuppression during postpartum mammary gland involution. (A) Growth of E0771 tumors injected into intact mammary fat pads of C57/BL6 wild type (WT) (*n*=22 tumors) or SEMA7A knockout (KO) (*n*=16 tumors) mouse one day post wean or involution day 1. Statistics are from day 13 tumor volumes using unpaired two-tailed Welch’s t test. (B) Growth of E0771 tumors injected into intact mammary fat pads of KO or SEMA7A heterozygous (Het) mouse mammary tissues of nulliparous or at involution day 1. Het nulliparous *n*=10 tumors; Het involution *n*=8 tumors; KO nulliparous *n*=8 tumors; KO involution *n*=6 tumors. Statistics are from day 16 tumor volumes using ordinary one-way ANOVA with Tukey’s multiple comparisons test. (C) Percent LECs expressing PD-L1 from nulliparous (Nul) or involution (Inv) days 2, 3, 4, 6, 8, and fully regressed (Reg) WT and KO mouse mammary glands by flow cytometry. WT gland sample sizes: Nul *n*=5; Inv2 *n*=5; Inv3 *n*=4; Inv4 *n*=6; Inv6 *n*=4; Inv8 *n*=5; Reg *n*=3. KO gland sample sizes: Nul *n*=6; Inv2 *n*=4; Inv3 *n*=3; Inv4 *n*=6; Inv6 *n*=5; Inv8 *n*=3; Reg *n*=2. Ordinary two-way ANOVA with Šídák’s multiple comparison test. (D) Percent total macrophages expressing PD-L1 from Nul, Inv 2, 3, 4, 6, 8, and Reg WT and KO mouse mammary glands by flow cytometry. WT gland sample sizes: Nul *n*=6; Inv2 *n*=4; Inv3 *n*=5; Inv4 *n*=6; Inv6 *n*=4; Inv8 *n*=6; Reg *n*=6. KO gland sample sizes: Nul *n*=6; Inv2 *n*=4; Inv3 *n*=5; Inv4 *n*=6; Inv6 *n*=4; Inv8 *n*=5; Reg *n*=5. Ordinary two-way ANOVA with Šídák’s multiple comparison test. (E) Percent M2-like macrophages expressing PD-L1 from Nul, Inv 2, 3, 4, 6, 8, and Reg WT and KO mouse mammary glands by flow cytometry. WT gland sample sizes: Nul *n*=5; Inv2 *n*=6; Inv3 *n*=5; Inv4 *n*=6; Inv6 *n*=4; Inv8 *n*=6; Reg *n*=6. KO gland sample sizes: Nul *n*=6; Inv2 *n*=4; Inv3 *n*=5; Inv4 *n*=6; Inv6 *n*=4; Inv8 *n*=6; Reg *n*=6. Ordinary two-way ANOVA with Šídák’s multiple comparison test. (F) Percent PoEMs expressing PD-L1 from Nul, Inv 2, 3, 4, 6, 8, and Reg WT and KO mouse mammary glands by flow cytometry. WT gland sample sizes: Nul *n*=3; Inv2 *n*=6; Inv3 *n*=5; Inv4 *n*=6; Inv6 *n*=4; Inv8 *n*=6; Reg *n*=6. KO gland sample sizes: Nul *n*=5; Inv2 *n*=4; Inv3 *n*=4; Inv4 *n*=6; Inv6 *n*=4; Inv8 *n*=6; Reg *n*=6. Ordinary two-way ANOVA with Šídák’s multiple comparison test. Error bars are mean ± SEM. *\*p<0.05, **p<0.01, ***p<0.001, ****p<0.0001*.

### SEMA7A and PD-L1 expression in cells of the TME

To determine whether similar phenotypes were observed in a model of SEMA7A+ BC, E0771 mouse mammary tumor cells engineered to overexpress (OE) SEMA7A were injected into nulliparous mice. Consistent with previously published studies^20,46,47^, SEMA7A OE tumors exhibited increased surface SEMA7A expression (observed by flow cytometry) and grew faster than empty vector controls (Ctrl) (Fig. 2A&B, SFig. 2A). The increased growth was accompanied by increases in surface expression of PD-L1 on the tumor cells (Fig. 2C), the CD45+ cells (Fig. 2D), the macrophages (F4/80+) (Fig. 2E), the M1-like macrophages (CD11b+/F4-80+/MHCII+/CD206-), the M2-like macrophages (CD11b+/F4-80+/MHCII-/CD206+), and the PoEMs (CD11b+/F4-80+/PDPN+) (Fig. 2F). Increased frequency of total and PD-L1+ LECs was also observed (SFig. 2B). Additionally, the observed increases in PD-L1+ immune cell populations were not due to increased total immune cell infiltration (SFig. 2C). However, a decrease in the frequency of M1-like macrophages that was accompanied by an increase in M2-like macrophages and PoEMs in the SEMA7A OE tumors was observed (SFig. 2D-G). Furthermore, decreases in the percentages of CD4+ and CD8+ T cells (Fig. 2G) alongside an increased PD-1 expression was observed in the OE group tumors (Fig. 2H). Finally, the T cells present in SEMA7A OE tumors exhibited markers of increased exhaustion (PD-1^hi^/LAG3^hi^/TNFα-/IFNγ-) (Fig. 2I) and decreased activation signatures (PD-1^lo/mid^/LAG3-/TNFα+/IFNγ+) (Fig. 2J). Overall, these results indicate that SEMA7A expression in mammary tumors helps drive an immunosuppressive TME by enriching for M2-like macrophages and PoEMs and through promoting T cell exhaustion (Fig. 2K).

**Figure 2:**
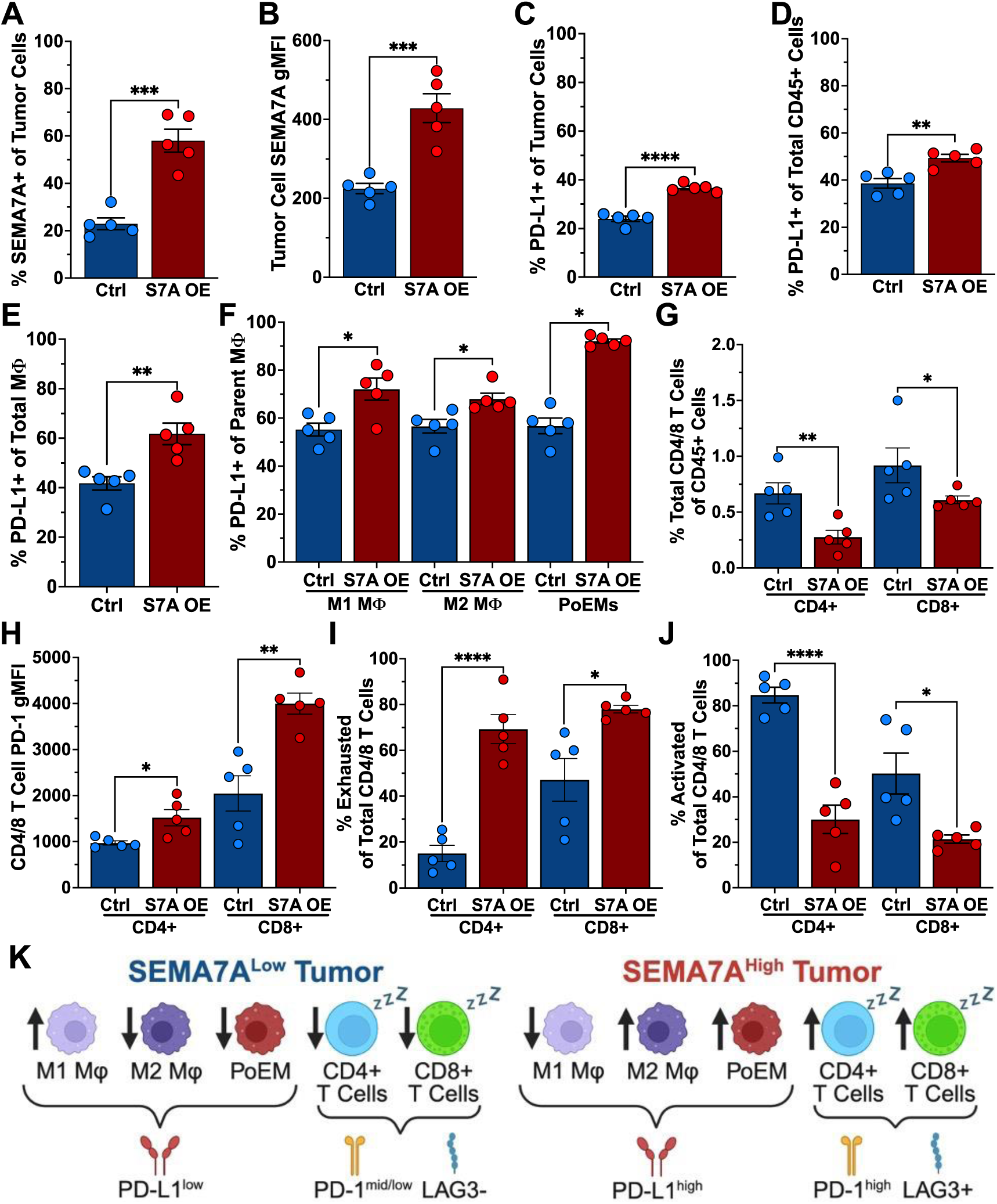
Tumor-derived SEMA7A in relation to PD-L1 and the tumor immune microenvironment. (A) Percent SEMA7A+ tumor cells isolated from E0771 control (Ctrl) and SEMA7A overexpressing (S7A OE) tumors. (B) Geometric mean fluorescence intensity (gMFI) of SEMA7A expression from Ctrl or S7A OE tumors. (C) Percent PD-L1+ tumor cells from Ctrl or S7A OE tumors. (D) Percent PD-L1+ immune cells from Ctrl or S7A OE tumors. (E) Percent PD-L1+ total macrophages from Ctrl or S7A OE tumors. (F) Percentage PD-L1+ M1-like macrophages, M2-like macrophages, and PoEMs from Ctrl or S7A OE tumors. (G) Frequency of CD4+ and CD8+ T cells of total immune cells from Ctrl or S7A OE tumors. (H) gMFI of PD-1 on CD4+ and CD8+ T cells from Ctrl or S7A OE tumors. (I) Percent exhausted (PD-1^high^/LAG-3+/IFNγ-/TNFα-) CD4+ and CD8+ T cells of total CD4/8 T cells from Ctrl or S7A OE tumors. (J) Percent activated (PD-1^mid^/LAG-3-/IFNγ+/TNFα+) CD4+ and CD8+ T cells of total CD4/8 T cells from Ctrl or S7A OE tumors. (K) Summary of macrophage and T cell phenotypes between SEMA7A^Low^ and SEMA7A^High^ tumors. Error bars are mean ± SEM. Sample sizes for A-J: Ctrl *n*=5 tumors; S7A OE *n*=5 tumors. Statistics for A-J are unpaired two-tailed Welch’s t tests. *\*p<0.05, **p<0.01, ***p<0.001, ****p<0.0001*.

Based on the enrichment for PD-L1+ tumor cells and cells of the TME in SEMA7A+ tumors, we investigated SEMA7A-induced signaling mechanisms that could result in upregulation of cell surface PD-L1 expression. To test this, we treated E0771 and 66cl4 mouse mammary carcinoma cells in vitro with exogenous purified SEMA7A and increased surface PD-L1 expression was observed by flow cytometry; further, expression of PD-L1 was abrogated by function blocking antibodies for integrin-α6/β1, a putative SEMA7A receptor. We also tested a novel SEMA7A blocking antibody, SmAbH1, and similarly observed reduced PD-L1 expression (SFig. 3A&B). SmAbH1 binds to SEMA7A (SFig. 3C) and reduces tumor cell viability in vitro (SFig. 3D). Specifically, using in vitro assays, inhibition of SEMA7A+ tumor cell growth and clonogenicity was observed with little to no effect in cells where SEMA7A was reduced via shRNA (KD) (SFig. 3E). When MDA-MB-231 breast cancer cells were similarly treated with exogenous SEMA7A, we confirmed increased PD-L1 expression by immunoblot, which was also decreased with the blocking antibodies (Fig. 3A). Previous studies identify several signaling cascades that could mediate increased expression of PD-L1 in response to SEMA7A ^46,47,50,68–75^ including integrin linked kinase (ILK) and PI3K, which can both result in activation of AKT via phosphorylation. In support of this potential signaling mechanism treatment with SEMA7A upregulated phosphorylation/activation of AKT, while receptor blockade reversed activation (Fig. 3B). There was also a modest increase in pSTAT3 at S727, which is associated with transcriptional regulation of PD-L1 expression though non-canonical STAT3 signaling^76–79^, which was also partially mitigated by receptor blockade (Fig. 3C). In support of a role for ILK and PI3K in initiating upregulation of PD-L1, targeted inhibitors of both kinases resulted in downregulation of pAKT and pSTAT3 (Fig. 3D&E) as well as PD-L1 expression. Furthermore, direct inhibition of STAT3 also resulted in downregulation of PD-L1 expression (Fig. 3F).

**Figure 3:**
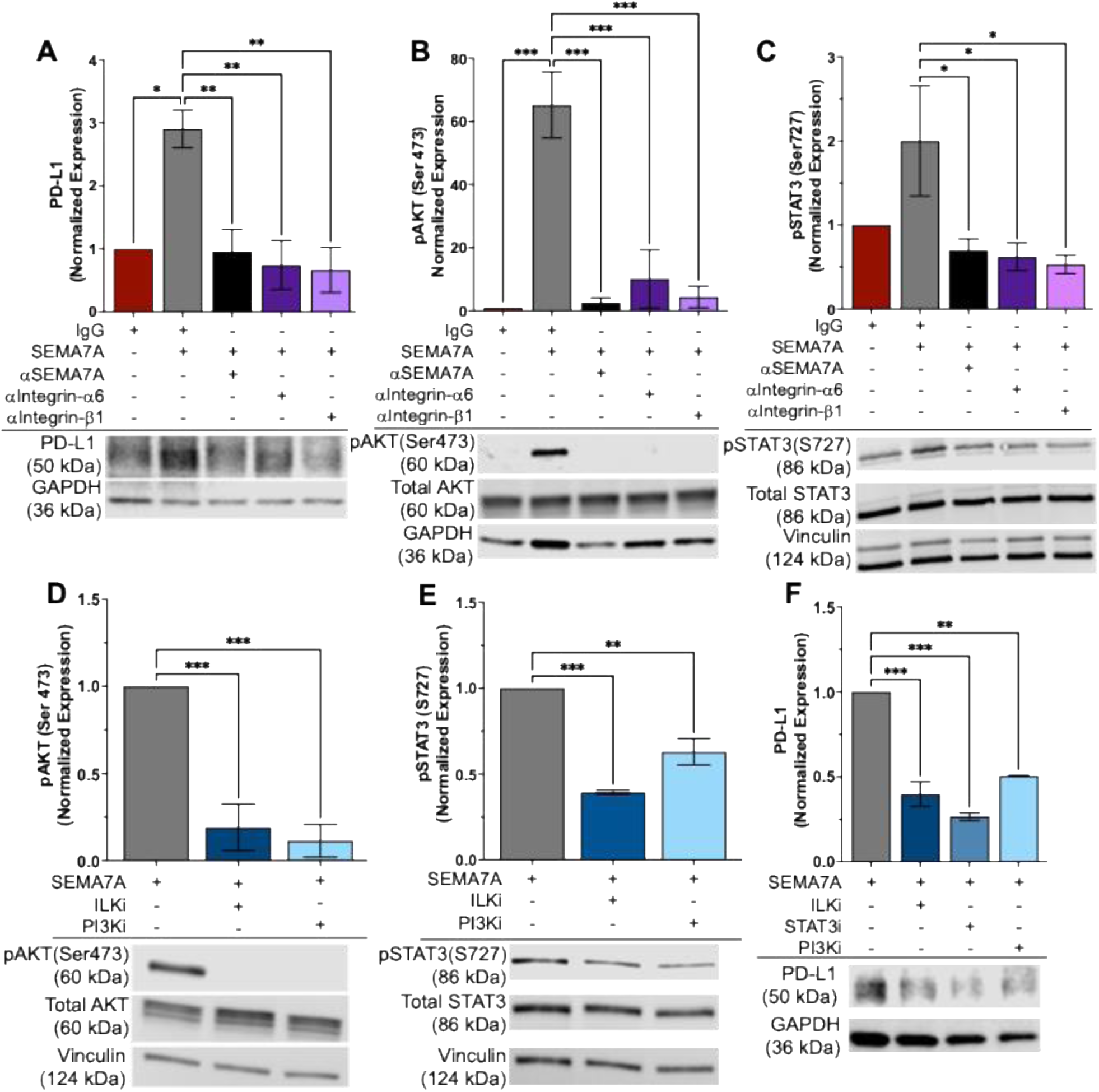
SEMA7A-mediated signaling and PD-L1 expression. (A) Immunoblot and densiometric quantification of PD-L1 protein expression in MDA-MB-231 human breast cancer cells following treatment with SEMA7A +/-inhibitors of SEMA7A, integrin-α6, or integrin-β1. (B) Immunoblot and densiometric quantification of pAKT and total AKT expression in MDA-MB-231 cells following treatment with SEMA7A +/-inhibitors of SEMA7A, integrin-α6, or integrin-β1. (C) Immunoblot and densiometric quantification of pGSK3β and total GSK3β expression in MDA-MB-231 cells following treatment with SEMA7A +/-inhibitors of SEMA7A, integrin-α6, or integrin-β1. (D) Immunoblot and densiometric quantification of pAKT and total AKT expression in MDA-MB-231 cells following treatment with SEMA7A +/-inhibitors of ILK or PI3K. (E) Immunoblot and densiometric quantification of pGSK3β and total GSK3β expression in MDA-MB-231 cells following treatment with SEMA7A +/-inhibitors of ILK or PI3K. (F) Immunoblot and densiometric quantification of PD-L1 expression in MDA-MB-231 cells following treatment with SEMA7A +/-inhibitors of ILK or PI3K. (G) Left: Schematic depicting SEMA7A signaling through integrin-α6/β1 to activate ILK and/or AKT to phosphorylate GSK3β and allow for stabilization/accumulation of PD-L1 protein. Right: Schematic depicting inhibition of SEMA7A signaling results in active GSK3β, which then phosphorylates PD-L1 and targets it for ubiquitin-mediated protein degradation. Immunoblots shown are representative from 3 independent experiments and error bars on quantification are mean ± SD from 3 biologic replicates. Statistics on densiometric quantification are ordinary one-way ANOVA with Tukey’s multiple comparisons test. *\*p<0.05, **p<0.01, ***p<0.001, ****p<0.0001*.

### Immune checkpoint inhibition and SEMA7A in preclinical models of breast cancer

To test whether immune checkpoint inhibiting therapies could decrease growth and alter the immunosuppressive characteristics of SEMA7A OE tumors we treated with anti-PD-1 and anti-PD-L1 (αPD-1/αPD-L1) directed monoclonal antibodies or IgG controls. Initially, slower tumor growth rates were observed in the SEMA7A OE tumors, which was followed by a rebound in tumor growth. Conversely, tumor growth was not slowed in the Ctrl tumors with treatment (Fig. 4A&B). Flow cytometric analysis revealed an enrichment for SEMA7A expressing cells with treatment as well as higher overall levels of SEMA7A expression at study end (Fig. 4C&D). Reduction in PD-L1+ tumor cells was observed in the OE tumors that were treated with αPD-1/αPD-L1, which was not observed in the Ctrl tumors (Fig. 4E). Further analysis of the Ctrl and SEMA7A OE groups for PD-L1+ cell populations in the TME, (described in Figure 2) revealed increased PD-L1+ total immune cells in both treated and untreated groups (SFig. 4A) as well as small, but significant, decreases in PD-L1+ LECs, total and M1-like macrophages, and PoEMs that were exclusive to the SEMA7A OE tumors (SFig. 4B-F). Additionally, we observed that treatment promoted infiltration of total and M1 macrophages (Fig. 4F&G), with little to no change in M2 macrophages (Fig. 4H), and significantly decreased PoEM presence in the OE tumors (Fig. 4I), which was not observed in the Ctrl tumors (Fig. 4H&I).

**Figure 4:**
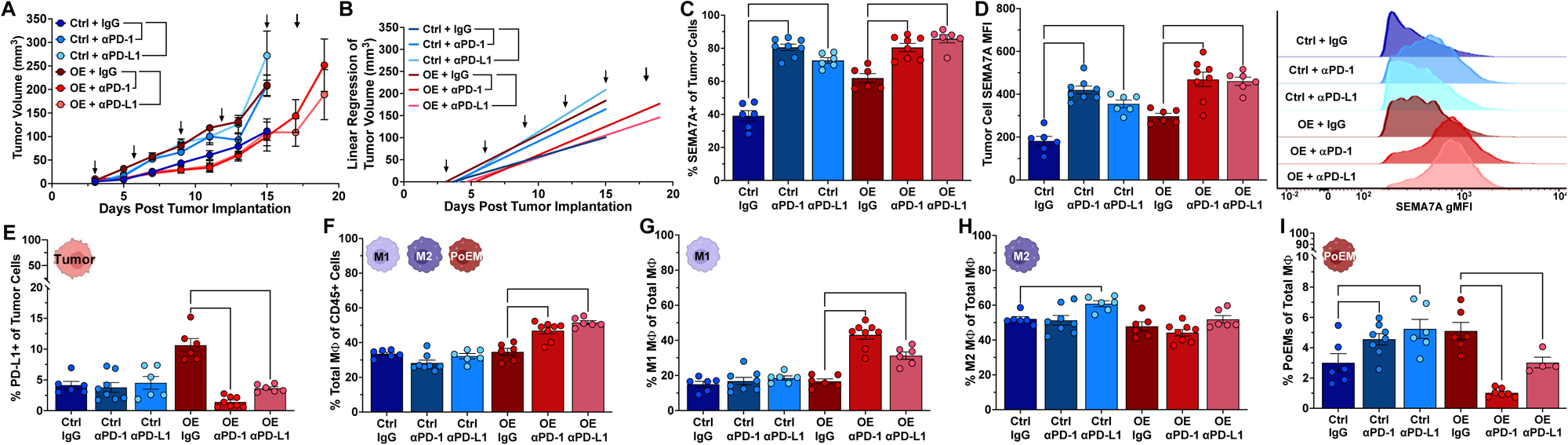
SEMA7A tumors treated with αPD-1/αPD-L1 antibodies. (A) Tumor growth in E0771 control (Ctrl) or SEMA7A overexpressing (OE) tumors. Arrows indicate administration of treatment with IgG control, αPD-1, or αPD-L1. All Ctrl groups and OE + IgG were taken down at day 15 due to early signs of ulceration. OE + αPD-1 and OE + αPD-L1 were taken down at day 20 due to early signs of ulceration. Statistics shown are from day 15 tumor volumes using ordinary one-way ANOVA with Tukey’s multiple comparisons test. (B) Growth rates from (A). (C) Percent SEMA7A+ tumor cells from Ctrl or OE tumors following treatments outlined in (A). (D) Median fluorescence intensity (MFI) of SEMA7A expression on tumor cells from Ctrl or OE tumors following treatments outlined in (A). Quantification (left) and representative histograms (right). (E) Percent PD-L1+ tumor cells from Ctrl or OE tumors following treatments outlined in (A). (F) Percent M1-like macrophages from Ctrl or OE tumors following treatments outlined in (A). (G) Percent M2-like macrophages from Ctrl or OE tumors following treatments outlined in (A). (H) Percent PoEMs of total macrophages from Ctrl or OE tumors following treatments outlined in (A). (I) Percent macrophages of total immune cells from Ctrl or OE tumors following treatments outlined in (A). Error bars as mean ± SEM. Sample sizes for A-I: Ctrl + IgG *n*=6 tumors; Ctrl + αPD-1 *n*=8 tumors; Ctrl + αPD-L1 *n*=6 tumors; OE + IgG *n*=6 tumors; OE + αPD-1 *n*=8 tumors; OE + αPD-L1 *n*=6 tumors. Statistics for C-I are ordinary one-way ANOVA with Tukey’s multiple comparisons test. *\*p<0.05, **p<0.01, ***p<0.001, ****p<0.0001*.

Treated SEMA7A OE tumors also displayed increased frequency of both CD4+ and CD8+ T cells (Fig. 5A&B) and expression levels of PD-1 on T cells was reduced by αPD-1 treatment (Fig. 5C&D), which indicates a partial reversion of exhaustion. Additionally, in OE treated tumors, we observed decreased exhausted and increased activated CD8+ T cell frequency with both treatments and αPD-1 also reduced exhausted and increased activated CD4+ T cells (Fig. 5E-H), all of which did not occur in Ctrl tumors. Utilizing another model of TNBC, the effect of SEMA7A overexpression on tumor cell PD-L1 expression, as well as the immunosuppressive effect of SEMA7A on the composition of the immune TME was validated (SFig. 5A-T).

**Figure 5:**
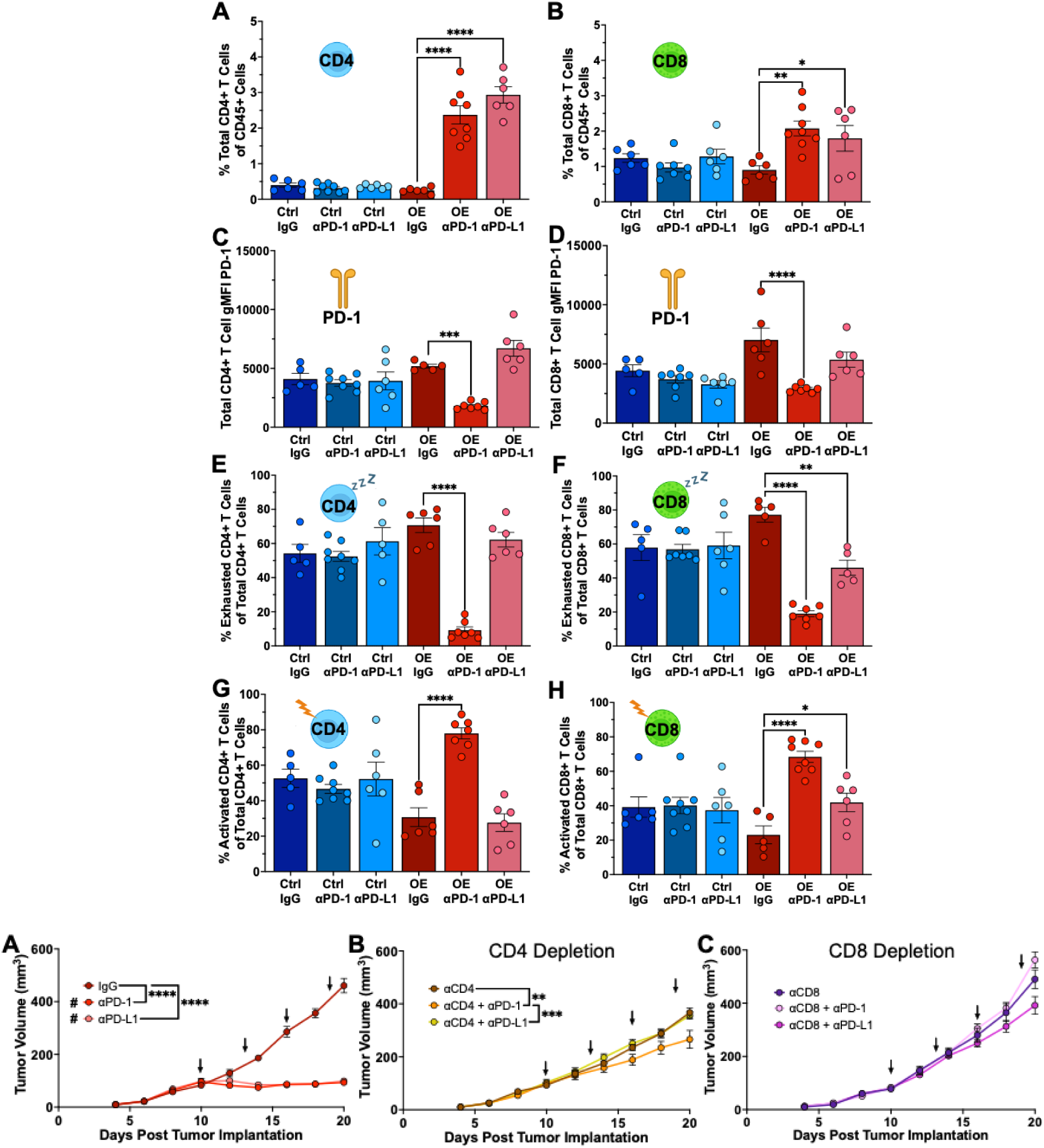
αPD-1/αPD-L1 antibodies and T cell infiltration and activation in SEMA7A OE tumors. (A) Percent CD4+ T cells from E0771 control (Ctrl) or SEMA7A overexpressing (OE) tumors following treatment with IgG control, αPD-1, or αPD-L1. (B) Percent CD8+ T cells from Ctrl or OE tumors following treatment with IgG control, αPD-1, or αPD-L1. (C) PD-1 gMFI on CD4+ T cells from Ctrl or S7A OE tumors following treatment with IgG control, αPD-1, or αPD-L1. (D) PD-1gMFI on CD8+ T cells from Ctrl or S7A OE tumors following treatment with IgG control, αPD-1, or αPD-L1. (E) Percent exhausted (PD-1^high^/LAG-3+/IFNγ-/TNFα-) CD4+ T cells from Ctrl or S7A OE tumors following treatment with IgG control, αPD-1, or αPD-L1. (F) Percent exhausted (PD-1^high^/LAG-3+/IFNγ-/TNFα-) CD8+ T cells from Ctrl or S7A OE tumors following treatment with IgG control, αPD-1, or αPD-L1. (G) Percent activated (PD-1^mid^/LAG-3-/IFNγ+/TNFα+) CD4+ T cells from Ctrl or S7A OE tumors following treatment with IgG control, αPD-1, or αPD-L1. (H) Percent activated (PD-1^mid^/LAG-3-/IFNγ+/TNFα+) CD8+ T cells from Ctrl or S7A OE tumors following treatment with IgG control, αPD-1, or αPD-L1. Error bars are mean ± SEM. Sample sizes for A-H: Ctrl + IgG *n*=6 tumors; Ctrl + αPD-1 *n*=8 tumors; Ctrl + αPD-L1 *n*=6 tumors; OE + IgG *n*=6 tumors; OE + αPD-1 *n*=8 tumors; OE + αPD-L1 *n*=6 tumors. Statistics for A-H are ordinary one-way ANOVA with Tukey’s multiple comparisons test. *\*p<0.05, **p<0.01, ***p<0.001, ****p<0.0001*.

Finally, to show that the effect of αPD-1/αPD-L1 on SEMA7A OE tumor growth is reliant on T cells, we depleted T cells using αCD4 or αCD8 monoclonal antibodies followed by treatment with αPD-1/PD-L1. Consistent with previous results, treatment significantly decreased tumor growth (Fig. 6A, STable 1, and SFig. 6A&B). However, depletion of CD4+ and CD8+ T cells blocked the effects of the treatments on tumor growth (Fig. 6B&C, SFig. 6C&D), indicating that αPD-1/αPD-L1 based treatments are reliant on both CD4+ and CD8+ T cells in SEMA7A OE tumors.

**Figure 6:**
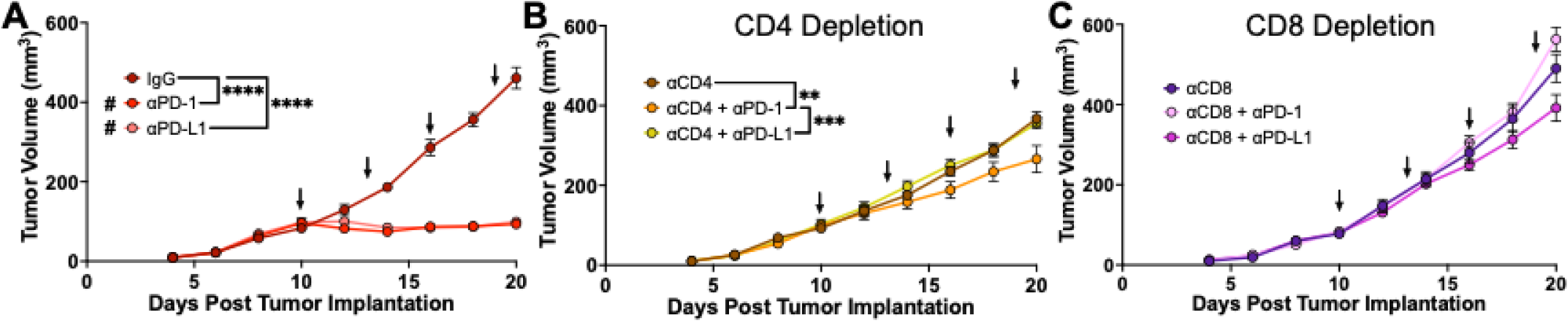
αPD-1/αPD-L1 efficacy in SEMA7A OE tumors with depletion of CD4+ and CD8+ T cells. (A) Growth of SEMA7A overexpressing E0771 tumors. Arrows indicate administration of treatment with IgG control, αPD-1, or αPD-L1. IgG *n*=6 tumors; αPD-1 *n*=6 tumors; αPD-L1 *n*=8 tumors. (B) Growth of SEMA7A overexpressing E0771 tumors in CD4 T cell-depleted mice with/without αPD-1/αPD-L1 treatment. αCD4 was administered at days 6, 8, 13, and 18. Arrows represent IgG, αPD-1, or αPD-L1 treatments. αCD4 + IgG *n*=6 tumors; αCD4 + αPD-1 *n*=6 tumors; αCD4 + αPD-L1 *n*=8 tumors. (C) Growth of SEMA7A overexpressing E0771 tumors in CD8 T cell-depleted mice with/without αPD-1/αPD-L1 treatment. αCD8 was administered at days 6, 8, 13, and 18. Arrows represent IgG, αPD-1, or αPD-L1 treatments. αCD8 + IgG *n*=6 tumors; αCD8 + αPD-1 *n*=6 tumors; αCD8 + αPD-L1 *n*=8 tumors. Error bars are mean ± SEM. Statistics are from day 20 using ordinary one-way ANOVA with Tukey’s multiple comparisons test. *\*p<0.05, **p<0.01, ***p<0.001, ****p<0.0001.* # indicates a statistically significant difference from all other groups (Supplemental Table 1).

### Direct targeting of SEMA7A and anti-tumor immune responses

Next, a novel SEMA7A mouse monoclonal antibody-based treatment was compared to αPD-1/αPD-L1 in preclinical murine models of TNBC. SmAbH1 treatment significantly decreased tumor growth and extended mouse survival probability in vivo (Fig. 7A) as measured by time to maximal tumor volume/burden or moribund criteria (Fig. 7B); additionally, 60% of tumors completely regressed in the treated group with the other 40% demonstrating partial responses compared to no responses in the IgG controls (Fig. 7C). In a second model, SmAbH1 and αPD-L1 were directly compared to reveal significantly decreased tumor growth in the SmAbH1 group compared to αPD-L1 or IgG control (Fig. 7D&E). Additionally, targeting SEMA7A directly in this model resulted in a 70% complete response rate with the remaining 30% having partial responses; comparatively, αPD-L1 had a 73% partial response rate with no complete responders (Fig. 7F).

**Figure 7:**
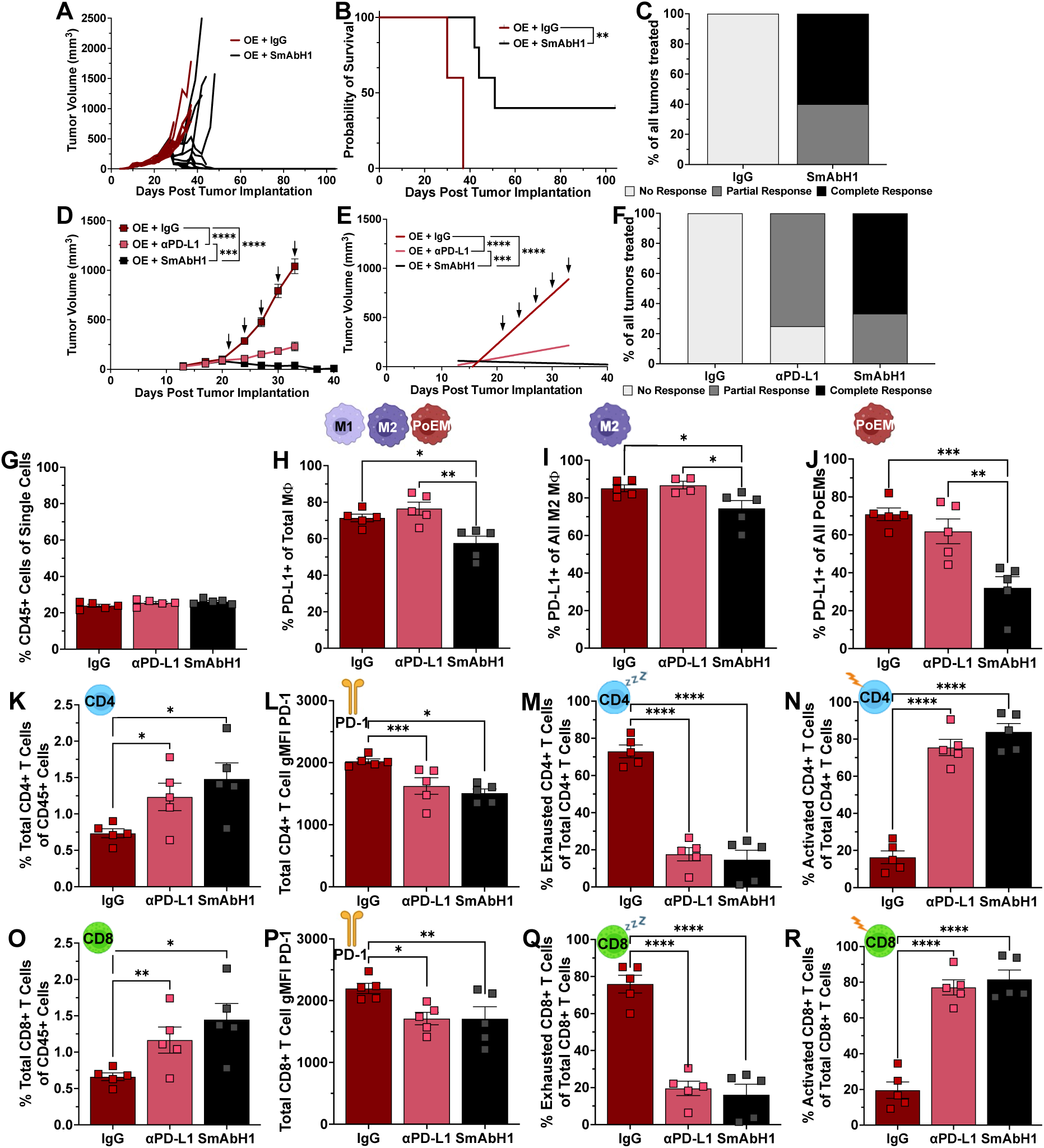
Direct targeting of SEMA7A in vivo. (A) Growth of individual E0771 SEMA7A overexpressing (OE) tumors following treatment with IgG control (*n*=10 tumors) or αSEMA7A (SmAbH1) (*n*=10 tumors). (B) Survival analysis from (A). *n*=5 mice/group with Mantel-Cox test. (C) Response rates of individual tumors from (A). *n*=10 tumors/group. (D) Growth of 66cl4 OE tumors following treatment with IgG control (*n*=12 tumors), SmAbH1 (*n*=12 tumors), or aPD-L1 (*n*=12 tumors). 6/6 OE + IgG and 6/6 OE + aPD-L1 mice were euthanized at day 33 due to tumor burden or morbidity, 3/6 OE + SmAbH1 mice were euthanized for time matched flow cytometry. Statistics are from day 33 tumor volumes using ordinary one-way ANOVA with Tukey’s multiple comparisons test. (E) Growth rates from (D). (F) Response rates of individual tumors from (D). *n*=12 tumors/group. (G) Percent immune cells from tumors following treatments outlined in (D). (H) Percent PD-L1+ total macrophages from tumors following treatments outlined in (D). (I) Percent PD-L1+ M2-like macrophages from tumors following treatments outlined in (D). (J) Percent PD-L1+ PoEMs from tumors following treatments outlined in (D). (K) Percent CD4+ T cells from tumors following treatments outlined in (D). (L) PD-1 geometric mean fluorescence intensity (gMFI) on CD4+ T cells from tumors following treatments outlined in (D). (M) Percent exhausted (PD-1^high^/LAG-3+/IFNγ-/TNFα-) CD4+ T cells from tumors following treatments outlined in (D). (N) Percent activated (PD-1^mid^/LAG-3-/IFNγ+/TNFα+) CD4+ T cells from tumors following treatments outlined in (D). (O) Percent CD8+ T cells from tumors following treatments outlined in (D). (P) PD-1 gMFI on CD8+ T cells from tumors following treatments outlined in (D). (Q) Percent exhausted (PD-1^high^/LAG-3+/IFNγ-/TNFα-) CD8+ T cells from tumors following treatments outlined in (D). (R) Percent activated (PD-1^mid^/LAG-3-/IFNγ+/TNFα+) CD8+ T cells from tumors following treatments outlined in (D). Error bars are mean ± SEM. Sample sizes for G-R: *n*=5 tumors/group. Statistics for G-R are ordinary one-way ANOVA with Tukey’s multiple comparisons test. *\*p<0.05, **p<0.01, ***p<0.001, ****p<0.0001*.

To determine whether SmAbH1 also reactivates anti-tumor immunity, tumors were harvested for flow cytometry. While there was no change in overall immune cell presence with SmAbH1 treatment, increased presence of M1-like macrophages, decreased presence of total, M2-like, and PoEMs as well as decreased PD-L1+ LECs and all PD-L1+ subtypes of macrophages was observed (Fig. 7G-J, SFig. 7A-F). Additionally, there were increased CD4+ and CD8+ T cells with decreased markers of exhaustion, which correlated with the increased presence of activated T cells (Fig.7 K-R). Collectively, these results indicate that SmabH1 may stimulate anti-tumor immunity, as well as tumor cell death, to promote regression of SEMA7A+ mammary tumors.

### SEMA7A and PD-L1 signaling in breast cancer patients

To determine if SEMA7A expression correlates with expression of markers associated with immunosuppression in breast cancer patient datasets, we analyzed The Cancer Genome Atlas (TCGA) breast cancer dataset using the Xena Functional Genomics Explorer^80^ for co-expression of *SEMA7A* mRNA with mRNAs for PD-1/PD-L1 (*PDCD1* and *CD274*, respectively), podoplanin (*PDPN*), *CD68* for general macrophages, and CD206 (or mannose receptor, *MRC1*) for M2 macrophages. In SEMA7A^high^ tumors, we observed significantly increased expression in comparison to SEMA7A^low^ tumors (Fig. 8A). Then, using the TIMER tool^81^, we performed correlation analysis of *SEMA7A* expression with *PDCD1, CD274, PDPN, CD68, MRC1, ILK,* and *STAT3* across all breast cancers as well as with stratified by breast cancer subtypes (i.e., Basal, Her2, Luminal A, and Luminal B). We identified that *SEMA7A* significantly correlated with each gene of interest across all breast cancers regardless of subtype with the exception of ILK and STAT3, which are primarily regulated at the protein level (Fig. 8B, SFig. 8A&B). KM Plotter^82^ analysis also revealed that overall survival is decreased in breast cancer patients with co-expression of *SEMA7A*, *CD274* (HR=1.36, p=0.023) (Fig. 8C) and *PDCD1* (HR=1.61, p=0.02) (Fig. 8D), which was not observed in patients with low expression of SEMA7A, regardless of PD-L1 or PD-1 expression (SFig. 8C&D). Finally, using survival analysis tools on a pan-cancer immunotherapy treatment dataset, expression of SEMA7A significantly predicts for decreased overall survival in patients treated with immunotherapy (HR=1.98, p=4.5e-6) (Fig. 8E) with SEMA7A^high^ and SEMA7A^low^ survival rates remaining identical until ∼10 months when the overall survival rate of SEMA7A^high^ patients significantly decreased.

**Figure 8:**
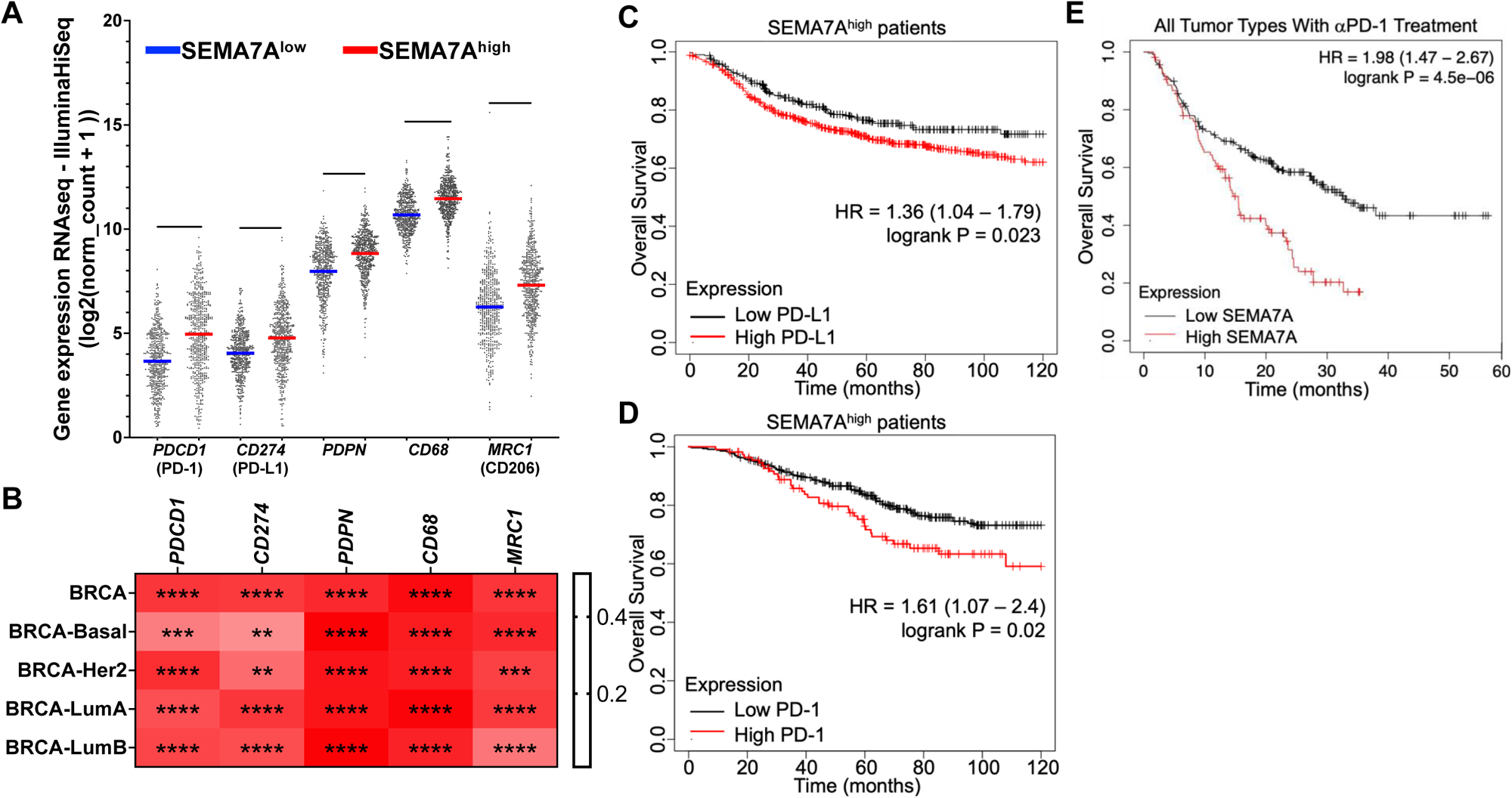
SEMA7A correlation with genes related to immunosuppression and patient survival analysis. (A) Co-expression analysis from The Cancer Genome Atlas (TCGA) breast cancer dataset using the Xena Functional Genomics Explorer^80^ for *SEMA7A, PDCD1*, *CD274*, *PDPN*, *CD68*, and *MRC1*. Unpaired two-tailed t test. (B) TIMER analysis^81^ from TCGA breast cancer data set for gene correlations between *SEMA7A* and *PDCD1*, *CD274*, *PDPN*, *CD68*, or *MRC1*. Gene correlations stratified by breast cancer subtypes. Heatmaps generated using purity-adjusted partial spearman’s rank correlation coefficient. (C) Overall survival analysis of SEMA7A^high^ breast cancer patients stratified by high or low expression of PD-L1 using the KM Plotter tool^82^. (D) Overall survival analysis of SEMA7A^high^ breast cancer patients stratified by high or low expression of PD-1 using the KM Plotter tool^82^. (E) Overall survival analysis of patients that received aPD-1 therapy in all cancers and stratified by high or low expression of SEMA7A using the KM Plotter tool^82^. *\*p<0.05, **p<0.01, ***p<0.001, ****p<0.0001*.

## Discussion

The remodeling of the postpartum mammary gland back to the pre-pregnant state after lactation is necessary to prepare the tissue for subsequent rounds of lactation, but also results in upregulation of multiple pro-tumor pathways^18,22,37,38,43,83^. Our results presented herein suggest that SEMA7A promotes postpartum tumor growth in mice. One mechanism suggested by our studies is that host derived SEMA7A promotes expression of PD-L1 in multiple cell types in the mammary gland during postpartum involution to support an immunosuppressive environment, which could then be maintained in the tumor microenvironment of PPBCs. While our models of involution and SEMA7A tumors are all in mouse mammary tumor models, which do not completely represent the heterogeneity of human breast cancer subtypes, our patient data suggests that the mechanisms uncovered by our models may be at play in breast cancer patients. Additionally, extending our data to all cancers treated with immunotherapy suggests that SEMA7A^hi^ patients are very likely to develop resistance to immunotherapy and disease progression will occur. In support of this, treatment with immune checkpoint inhibitors in our mouse models resulted in enrichment for SEMA7A^hi^ tumor cells. Thus, directly targeting SEMA7A in patients that don’t respond to immunotherapy could be a potential therapeutic option. In support of this, we reveal for the first time that direct targeting with an anti-SEMA7A monoclonal antibody causes tumor regression while also promoting an anti-tumor immune response in mouse models of TNBC.

We postulate that the cell surface SEMA7A expressed on numerous cell types in the mammary gland during involution can be shed and signal to the implanted tumor cells which could result in activation of cell survival signaling and PD-L1 on the tumor cells. This is supported by our studies linking SEMA7A signaling to downstream activation of pro-survival kinases, ILK and AKT that have known functions in tumor progression including growth, invasion and metastasis. Furthermore, given the numerous parallels between the cells of the tissue microenvironment present during mammary gland involution and immune checkpoint signaling, we suggest that SEMA7A could further stimulate tumor growth by promoting immune evasion^17,22,23,27,38,41–44,84^. For example, during normal homeostatic conditions, PD-L1 binds to PD-1 on T cells to inhibit self-reactive T cells and to disrupt overreactive immune responses^85^. This mechanism may be crucial during postpartum mammary gland involution where death of the secretory mammary epithelial cells could result in presentation of self-and/or neo-antigens, such as milk proteins, that the immune system has not previously encountered^26,49,86–89^. Such a scenario could require activation of immune tolerance, which is supported by our evidence that harvested mammary glands from SEMA7A KO mice exhibit fewer immunosuppressive cells than the WT controls. However, we do not observe autoimmunity suggesting that other aspects of tolerance are involved. Additionally, we have observed that SEMA7A-driven traits are durably maintained in the tumor cells exposed to involution, which could allow them to travel freely throughout the lymph and blood vasculature and seed metastasis while avoiding immune destruction. These results may help explain why PPBC patients have higher rates of lymph node involvement, distant metastasis, and worse overall survival.

We explored the relationship between SEMA7A and PD-L1 to reveal that blockade of identified integrin receptors for SEMA7A and function blocking of SEMA7A can reduce PD-L1 expression. One potential explanation is through activation of AKT, which is supported by our results showing that inhibition of PI3K and ILK, both upstream of AKT, resulted in decreased SEMA7A induction of PD-L1 expression. Still, we did not observe a complete reduction in PD-L1, indicating that there are multiple, functionally redundant, signaling pathways that can lead to upregulation of PD-L1. One such mechanism revealed by our studies is non-canonical STAT3 signaling; however, it is also possible that SEMA7A signaling may also stimulate secretion of cytokines that induce canonical STAT3 signaling via JAK activation. Previous studies have shown that SEMA7A signaling can induce the production and secretion of IL-6^90,91^. Therefore, it is possible that SEMA7A creates a complex signaling pathway that results in IL-6 production, which then activates canonical JAK/STAT3 signaling, and finally transcriptional upregulation of PD-L1. Future studies are needed to further dissect the mechanism by which SEMA7A signaling results in activation of STAT3 via Y705 phosphorylation. An additional mechanism may be through AKT phosphorylation and inactivation of GSK3β, which has been shown to result in stabilization of the PD-L1 protein and may be an additional mechanism leading to the increase we observe^75^. Finally, AKT and ILK are also upstream of NF-κB, which is another known transcription factor that drives PD-L1 expression^71^. Additional studies are needed to further dissect the exact mechanisms by which SEMA7A mediates PD-L1 expression. Furthermore, co-targeting of multiple pro-survival signals with immunotherapy, such as those mediated by PI3K, ILK, GSK3β, or NF-κB could be explored in patients with SEMA7A+ breast cancers. In support of this, inhibitors of the GSK3β pathway are currently being tested as a stand-alone therapy and in combination with immunotherapy in multiple clinical trials^1,92,93^ and PI3K is also being targeted in hormone receptor positive cancers where we have shown that SEMA7A promotes endocrine therapy resistance^47,94^.

One limitation of this study is the utilization of 2 mouse mammary tumor models of triple negative breast cancer; however, current studies that are ongoing in the lab aim to include additional models of TNBC as well as ER+ models. Furthermore, given the complexity and potential overlap between SEMA7A signaling and signaling mediated by Her2, as well as the current availability of targeted therapy for these patients, we have deliberately focused on Her2-models. What is evident from our studies is a potential role for SEMA7A mediated immunosuppression in PPBCs, which we previously demonstrated by increased PD-L1 positivity in PPBCs compared to tumors from age-matched nulliparous women. Additionally, administration of αPD-1 therapy during postpartum involution was effective in pre-clinical mouse models of PPBC^42^. However, our current results reveal that enrichment for SEMA7A^high^ tumor cells is likely to occur with αPD-1/PD-L1 therapy. This may explain, in part, why the efficacy of therapies targeting PD-1/PD-L1 signaling in breast cancer patients is not completely understood^95^. In other types of solid tumors, including melanoma and lung, the mutational burden of a tumor is suggested to mediate immune recognition of neoantigens, which can initiate anti-tumor immune responses. In breast cancer, the mutational burden, degree of immune cell infiltration, and the infiltrate composition are inconsistent between subtypes and within each subtype^64,96–98^. Thus, many studies have attempted to identify clinically relevant neoantigens based off of predictive bioinformatic analyses and whole exome sequencing^97,99,100^. Unfortunately, transcription of the predicted neoantigens it is not guaranteed and many of the identified mutations are in proteins where clinical targeting has proved difficult based on significant treatment-related toxicities (e.g., PI3K inhibitors)^101,102^. While progress is being made on developing more specific drugs to target these and other neoantigens^103^, the ability for breast cancer cells to evade immune-mediated killing and removal still poses a great challenge in breast cancer. As such, we suggest that direct targeting of SEMA7A may have therapeutic advantage.

There are currently no targeted therapies directed at SEMA7A+ tumors and/or diagnostics for predicting SEMA7A status in breast cancer. Our results suggest that patients with SEMA7A+ cancers could partially benefit from αPD-1/αPD-L1 therapies, but that resistance may rapidly occur. Since significant adverse events are common and treatment efficacy is dependent on the patient’s ability to tolerate the treatment, it is possible that monitoring SEMA7A levels may be important for patients who receive immunotherapy^104–107^. Such monitoring could be accomplished by measuring blood and/or tissue levels of SEMA7A, as has been demonstrated in models of arthritis, ischemia, and in patients with myocardial infarction^91,108,109^. Additionally, as SEMA7A expression in adult tissues—outside of cancer—is limited to specific tissues and/or developmental/pathological events (e.g., mammary gland involution and inflammation) we suspect that SEMA7A-targeted therapy may have fewer off target side effects. Given that SEMA7A appears to be directly linked to tumor progression, our strategy for direct targeting of SEMA7A with a monoclonal antibody shows promise for future clinical trials with an optimized and humanized antibody that we are currently developing. An additional benefit of our antibody-based strategy would be the first targeted therapy for patients with PPBC— whose tumors appear to have a uniquely aggressive biology. Lastly, while we initially identified PPBC as a specific cancer type that may benefit from direct targeting of SEMA7A, we predict that that this therapeutic intervention would be beneficial for any patient with SEMA7A+ cancer. Thus, we will explore whether a companion diagnostic for detecting SEMA7A in tumors, like what is utilized Her2, has the potential to impact thousands of patients annually, as was observed with the discovery of Herceptin.

## Author contributions

Conception and design: A.E. and T.L.

Development of methodology: A.E., H.F., K.K., L.C., T.L.

Acquisition of data (provided animals, acquired and managed patients, provided facilities, etc.): A.E., A.B., H.F., K.K., L.C., T.L.

Analysis and interpretation of data (e.g., statistical analysis, biostatistics, computational analysis): A.E., H.F., K.K., L.C., A.B., T.L.

Writing, review, and/or revision of the manuscript: A.E., H.F., K.K., L.C., T.L.

Administrative, technical, or material support (i.e., reporting or organizing data, constructing databases): A.E., H.F., K.K., L.C., A.B., T.L.

Study supervision: A.E. and T.L.

## Acknowledgments

This work was supported by: NIH/NCI R01 CA211696-01A1 (to T.L.), NIH/NICHD R01 HD108335-01A1 (to T.L.), ACS RSG-16-171-010CSM (to T.L.), ACS MBGI-21-108-01-MBG (to T.L.), CU DOM ASPIRE Program (to T.L.), CU SPARK/REACH NIH: U01HL152405 and OEIDIT/AIA: MA2021-2028 (to T.L.), Gates Grubstake Award (to T.L.), Department of Medicine Outstanding Early Career Scholars Award (to T.L.), Carpenter Family Gift Fund (to T.L.), MSTP T32 1T32GM149361, Colorado CCTSI TL1 TR002533 (to A.E.), NIH/NCI NRSA 1F31CA268825-01A (to A.E.), and NIH/NCI Predoctoral to Postdoctoral Fellow Transition Award 1F99CA284276-01 (to A.E.). The authors acknowledge the CU Anschutz Cancer Center Cell Technologies Shared Resource for SEMA7A protein and SmAbH1antibody production, the CU Anschutz Cancer Center Flow Cytometry Shared Resource for flow cytometer use and technical assistance, the Cancer Center Support Grant (P30CA046934), and the Skin Diseases Research Cores Grant (P30AR057212). Contents are the authors’ sole responsibility and do not necessarily represent official NIH views. The authors would also like to thank the following persons for all their assistance with animal studies: Petra Dahms, Rachel Steinmetz, Alexander Stoller, Lyndsey Crump, Taylor Rutherford, and Sarah Tarullo. Lastly, the authors would like to thank the following persons for all their help in the production of this manuscript: Lyndsey Crump, Weston Porter, Rachel Carter, Senthil Muthuswamy, Ahn Lee, Steve Anderson, Eduardo Davila, Heide Ford, Jill Slansky, James DeGregori, Ross Kedl, and Virginia Borges. Graphical abstract and Figure 2K were made with BioRender.com.

## Supplemental figure titles and legends

**Figure S1:**
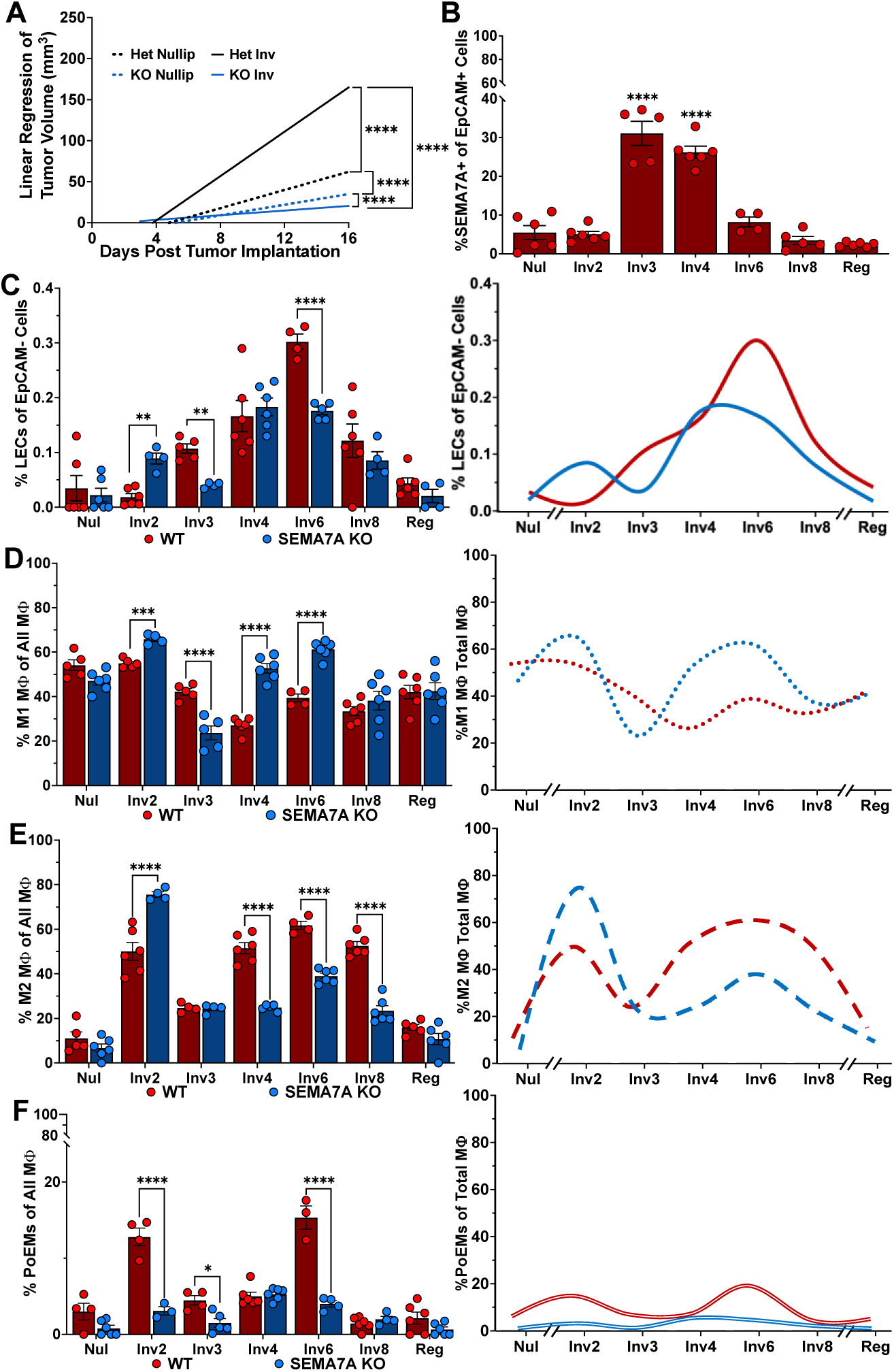
SEMA7A, LECs, and myeloid cells across mammary gland involution. (A) Growth rates of E0771 tumors injected into intact mammary fat pads of SEMA7A knockout (KO) or SEMA7A heterozygous (Het) mouse mammary tissues of nulliparous or at involution day 1. Het nulliparous *n*=10 tumors; Het involution *n*=8 tumors; KO nulliparous *n*=8 tumors; KO involution *n*=6 tumors. Statistics shown are from day 16 tumor volumes using ordinary one-way ANOVA with Tukey’s multiple comparisons test. (B) Percent SEMA7A+ EpCAM cells from nulliparous (Nul) or involution (Inv) days 2, 3, 4, 6, 8, and fully regressed (Reg) WT mouse mammary glands by flow cytometry. Mammary gland sample sizes: Nul *n*=6; Inv2 *n*=6; Inv3 *n*=5; Inv4 *n*=6; Inv6 *n*=4; Inv8 *n*=5; Reg *n*=6. Ordinary one-way ANOVA with Tukey’s multiple comparisons test. (C) Percent LECs from Nul, Inv 2, 3, 4, 6, 8, and Reg WT and KO mouse mammary glands by flow cytometry. WT glad sample sizes: Nul *n*=6; Inv2 *n*=6; Inv3 *n*=5; Inv4 *n*=6; Inv6 *n*=4; Inv8 *n*=6; Reg *n*=6. KO gland sample sizes: Nul *n*=6; Inv2 *n*=4; Inv3 *n*=4; Inv4 *n*=6; Inv6 *n*=5; Inv8 *n*=4; Reg *n*=4. (D) Percent M1-like macrophages from Nul, Inv 2, 3, 4, 6, 8, and Reg WT and KO mouse mammary glands by flow cytometry. WT gland sample sizes: Nul *n*=5; Inv2 *n*=5; Inv3 *n*=5; Inv4 *n*=6; Inv6 *n*=4; Inv8 *n*=6; Reg *n*=6. KO gland sample sizes: Nul *n*=6; Inv2 *n*=4; Inv3 *n*=5; Inv4 *n*=6; Inv6 *n*=6; Inv8 *n*=6; Reg *n*=6. (E) Percent M2-like macrophages expressing PD-L1 from Nul, Inv 2, 3, 4, 6, 8, and Reg WT and KO mouse mammary glands by flow cytometry. WT gland sample sizes: Nul *n*=5; Inv2 *n*=6; Inv3 *n*=4; Inv4 *n*=6; Inv6 *n*=4; Inv8 *n*=6; Reg *n*=5. KO gland sample sizes: Nul *n*=6; Inv2 *n*=4; Inv3 *n*=4; Inv4 *n*=5; Inv6 *n*=6; Inv8 *n*=6; Reg *n*=6. (F) Percent PoEMs from Nul, Inv 2, 3, 4, 6, 8, and Reg WT and KO mouse mammary glands by flow cytometry. WT gland sample sizes: Nul *n*=4; Inv2 *n*=4; Inv3 *n*=4; Inv4 *n*=6; Inv6 *n*=3; Inv8 *n*=6; Reg *n*=6. KO gland sample sizes: Nul *n*=6; Inv2 *n*=3; Inv3 *n*=5; Inv4 *n*=6; Inv6 *n*=4; Inv8 *n*=4; Reg *n*=6. Error bars are mean ± SEM. Statistics for C-F: ordinary two-way ANOVA with Šídák’s multiple comparison test. *\*p<0.05, **p<0.01, ***p<0.001, ****p<0.0001*.

**Figure S2:**
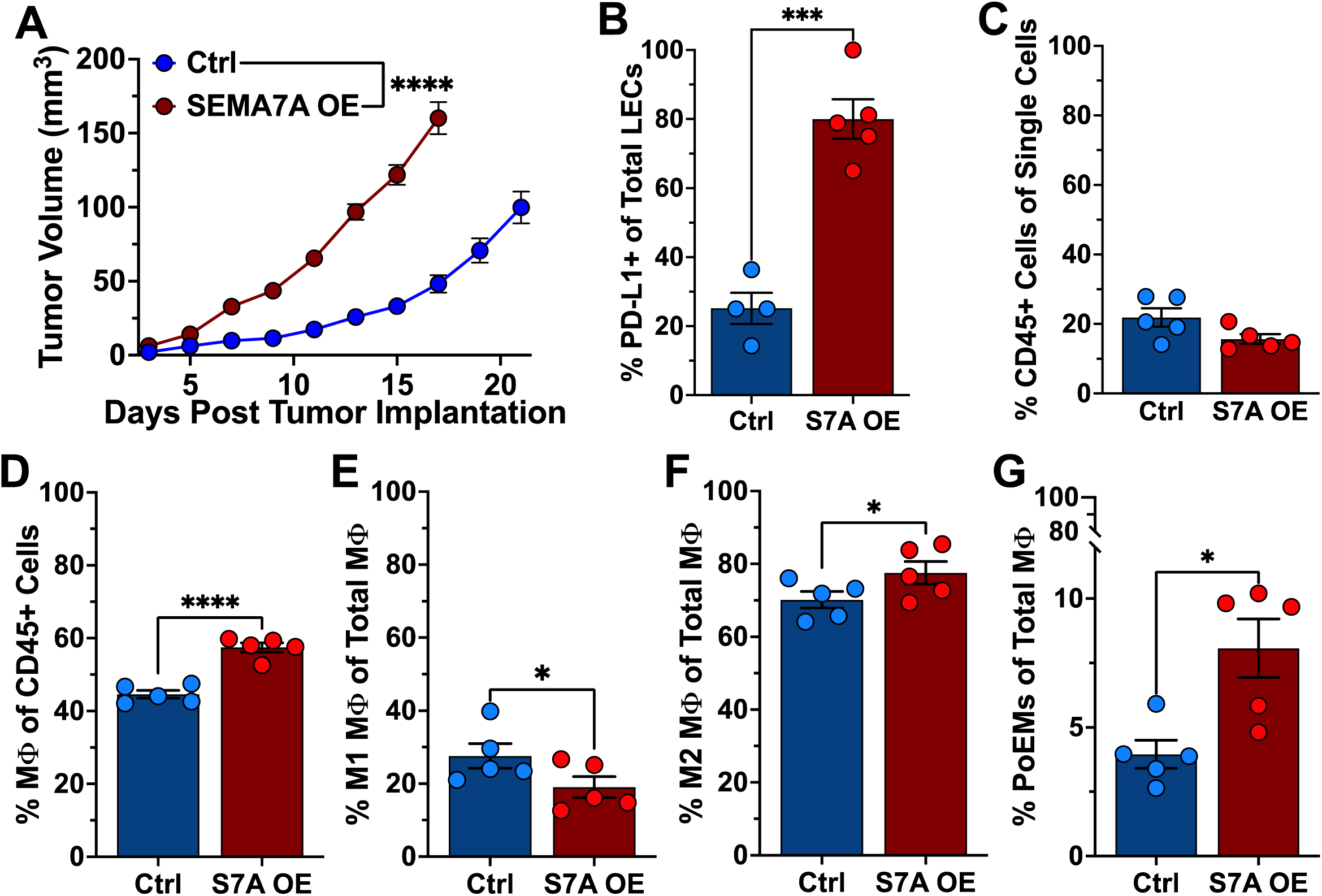
Tumor-derived SEMA7A and the tumor immune microenvironment. (A) Tumor growth in E0771 control (Ctrl) (*n*=10) or SEMA7A overexpressing (OE) (*n*=10) tumors. Statistics shown are from day 17 tumor volumes using unpaired two-tailed Welch’s t test. (B) Percent PD-L1+ LECs from Ctrl or S7A OE tumors from (A). (C) Percent immune cells from Ctrl or S7A OE tumors from (A). (D) Percent total macrophages from Ctrl or S7A OE tumors from (A). (E) Percent M1-like macrophages from Ctrl or S7A OE tumors from (A). (F) Percent M2-like macrophages from Ctrl or S7A OE tumors from (A). (G) Percent PoEMs from Ctrl or S7A OE tumors from (A). Error bars are mean ± SEM. Sample sizes for B-G: *n*=5 tumors/group. Statistics for B-G: unpaired two-tailed Welch’s t tests. *\*p<0.05, **p<0.01, ***p<0.001, ****p<0.0001*.

**Figure S3:**
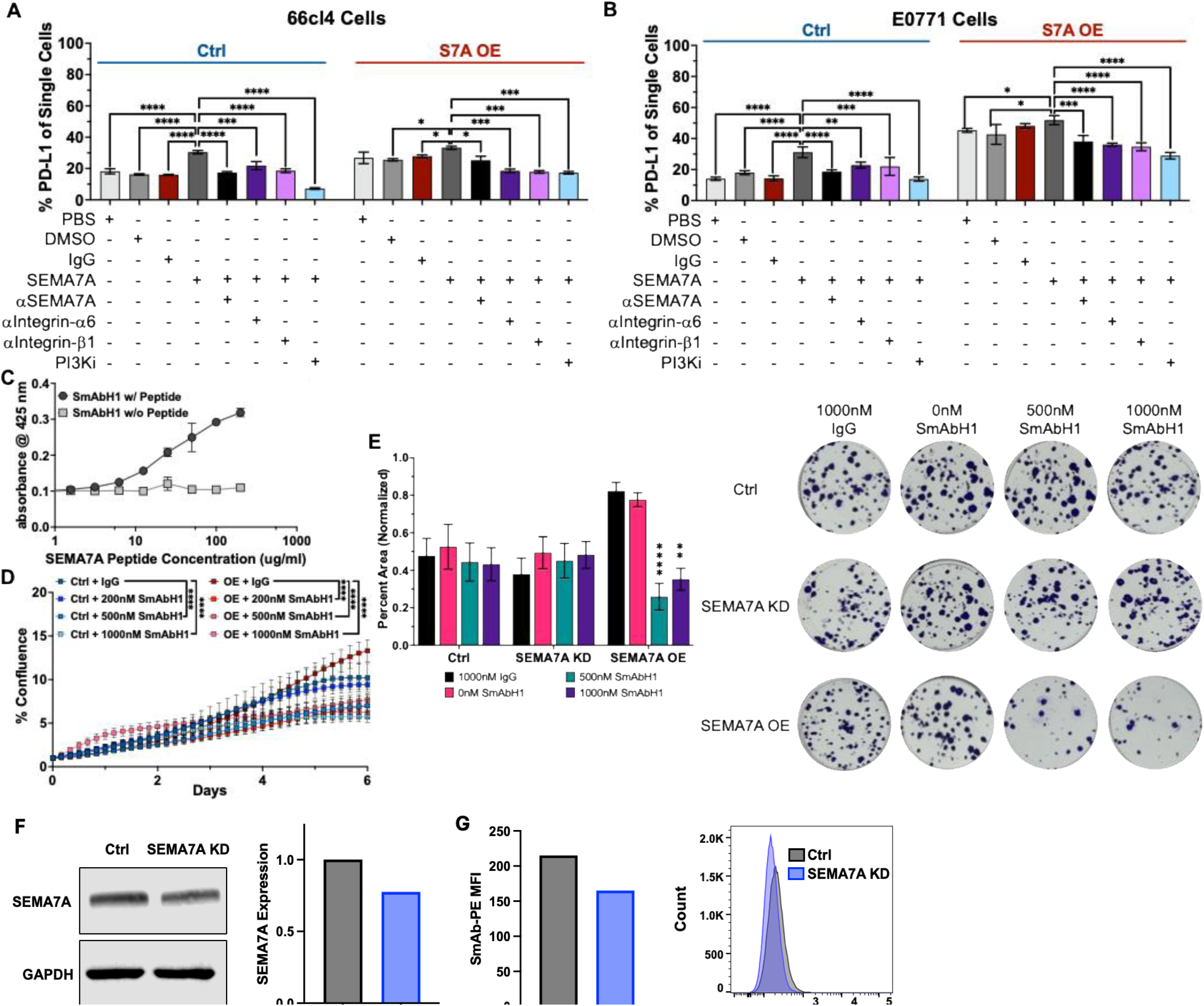
SEMA7A-mediated signaling and PD-L1 expression in mouse mammary carcinoma cells. (A) Percent PD-L1+ 66cl4 control (Ctrl) and SEMA7A overexpressing (S7A OE) cells following treatment with PBS, DMSO, or IgG for controls and SEMA7A +/-inhibitors of SEMA7A, Integrin-α6, Integrin-β1, or PI3K. (B) Percent PD-L1+ E0771 control (Ctrl) and SEMA7A overexpressing (S7A OE) cells following treatment with PBS, DMSO, or IgG for controls and SEMA7A +/-inhibitors of SEMA7A, Integrin-α6, Integrin-β1, or PI3K. Results are from pooled technical triplicates and error bars are mean ± SD and statistics shown are from ordinary two-way ANOVA with Šídák’s multiple comparison test. *\*p<0.05, **p<0.01, ***p<0.001, ****p<0.0001*.

**Figure S4:**
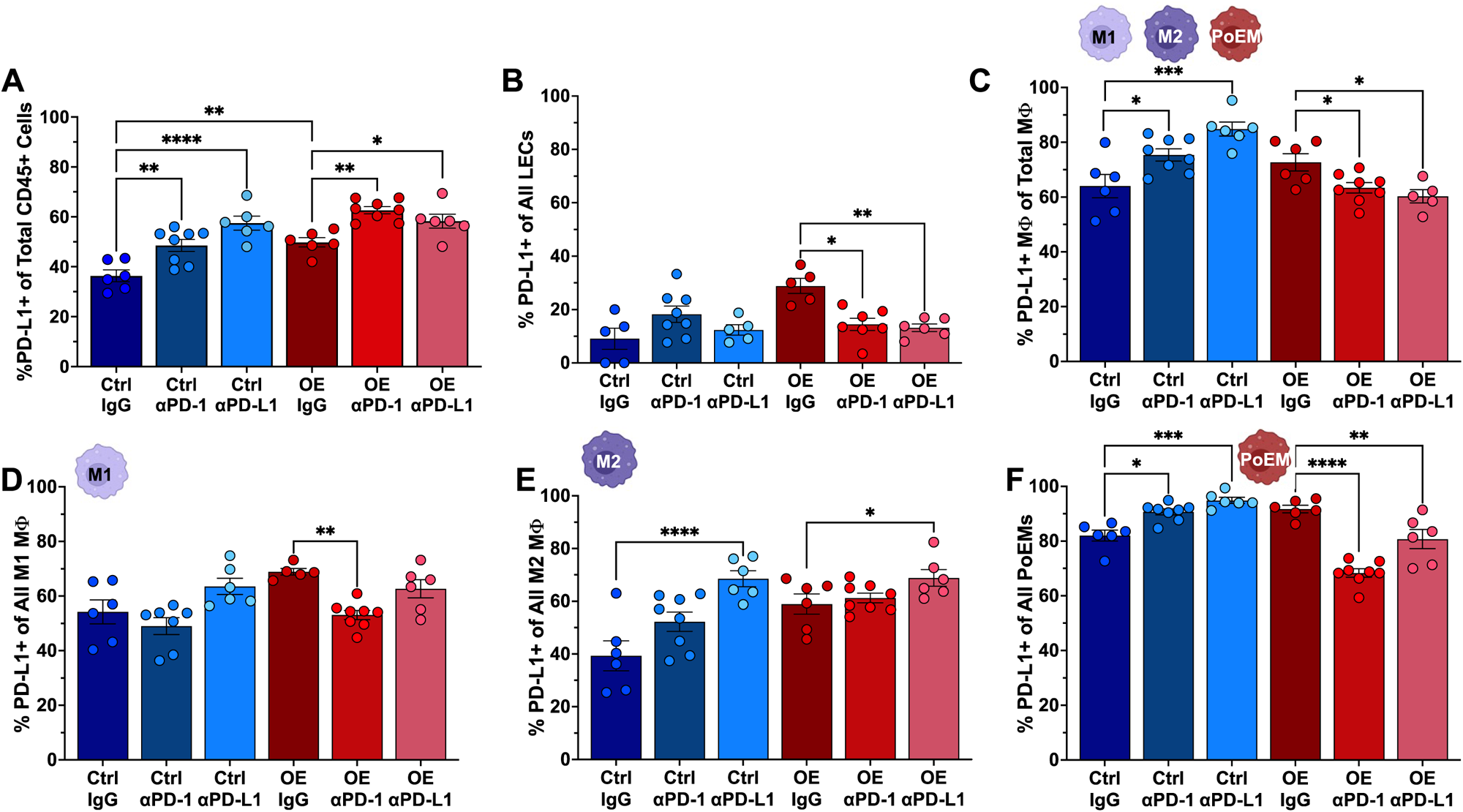
αPD-1/PD-L1 antibodies and the effect on PD-L1+ LECs and myeloid cells in SEMA7A tumors. (A) Percent PD-L1+ of total immune cells from control (Ctrl) or SEMA7A overexpressing (OE) tumors following treatments with IgG control, αPD-1, or αPD-L1. (B) Percent PD-L1+ LECs from Ctrl or OE tumors following treatments outlined in (A). (C) Percent PD-L1+ total macrophages from Ctrl or OE tumors following treatments outlined in (A). (D) Percent PD-L1+ M1-like macrophages from Ctrl or OE tumors following treatments outlined in (A). (E) Percent PD-L1+ M2-like macrophages from Ctrl or OE tumors following treatments outlined in (A). (F) Percent PD-L1+ PoEMs from Ctrl or OE tumors following treatments outlined in (A). Error bars are mean ± SEM. Sample sizes for A-F: Ctrl + IgG *n*=6 tumors; Ctrl + αPD-1 *n*=8 tumors; Ctrl + αPD-L1 *n*=6 tumors; OE + IgG *n*=6 tumors; OE + αPD-1 *n*=8 tumors; OE + αPD-L1 *n*=6 tumors. Statistics for A-F: ordinary one-way ANOVA with Tukey’s multiple comparisons test. *\*p<0.05, **p<0.01, ***p<0.001, ****p<0.0001*.

**Figure S5:**
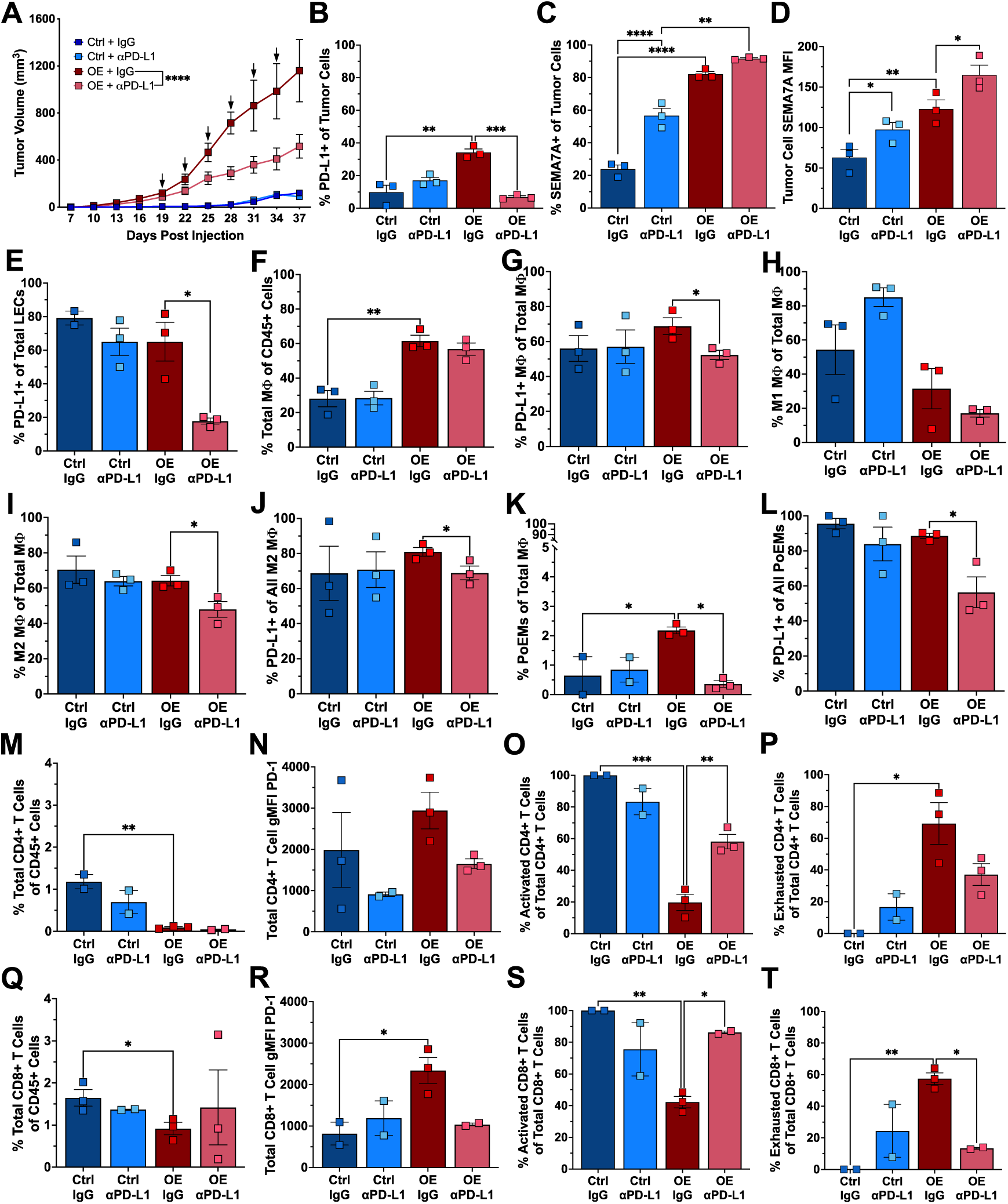
66cl4 SEMA7A tumors treated with **α**PD-L1. (A) Tumor growth in 66cl4 control (Ctrl) or SEMA7A overexpressing (OE) tumors. Arrows indicate administration of treatment with 250ug IgG control or αPD-L1. Ctrl + IgG *n*=6 tumors; Ctrl + αPD-L1 *n*=6 tumors; OE + IgG *n*=8 tumors; OE + αPD-L1 *n*=8 tumors. Statistics shown are from day 37 tumor volumes using ordinary one-way ANOVA with Tukey’s multiple comparisons test. (B) Percent PD-L1+ tumor cells from Ctrl or OE tumors following treatments outlined in (A). (C) Percent SEMA7A+ tumor cells from Ctrl or OE tumors following treatments outlined in (A). (D) Median fluorescence intensity (MFI) of SEMA7A expression on tumor cells from Ctrl or OE tumors following treatments outlined in (A). (E) Percent PD-L1+ LECs from Ctrl or OE tumors following treatments outlined in (A). (F) Percent total macrophages from Ctrl or OE tumors following treatments outlined in (A). (G) Percent PD-L1+ total macrophages from Ctrl or OE tumors following treatments outlined in (A). (H) Percent M1-like macrophages from Ctrl or OE tumors following treatments outlined in (A). (I) Percent M2-like macrophages from Ctrl or OE tumors following treatments outlined in (A). (J) Percent PD-L1+ M2-like macrophages from Ctrl or OE tumors following treatments outlined in (A). (K) Percent PoEMs from Ctrl or OE tumors following treatments outlined in (A). (L) Percent PD-L1+ PoEMs from Ctrl or OE tumors following treatments outlined in (A). (M) Percent CD4+ T cells from Ctrl or OE tumors following treatments outlined in (A). (N) PD-1 gMFI on CD4+ T cells from Ctrl or OE tumors following treatments outlined in (A). (O) Percent activated CD4+ T cells from Ctrl or OE tumors following treatments outlined in (A). (P) Percent exhausted CD4+ T cells from Ctrl or OE tumors following treatments outlined in (A). (Q) Percent CD8+ T cells from Ctrl or OE tumors following treatments outlined in (A). (R) PD-1 gMFI on CD8+ T cells from Ctrl or OE tumors following treatments outlined in (A). (S) Percent activated CD8+ T cells from Ctrl or OE tumors following treatments outlined in (A). (T) Percent exhausted CD8+ T cells from Ctrl or OE tumors following treatments outlined in (A). Error bars shown as mean ± SEM. Sample sizes for B-T: *n*=3 tumors/group. Statistics for B-T are ordinary one-way ANOVA with Tukey’s multiple comparisons test. *\*p<0.05, **p<0.01, ***p<0.001, ****p<0.0001*.

**Figure S6:**
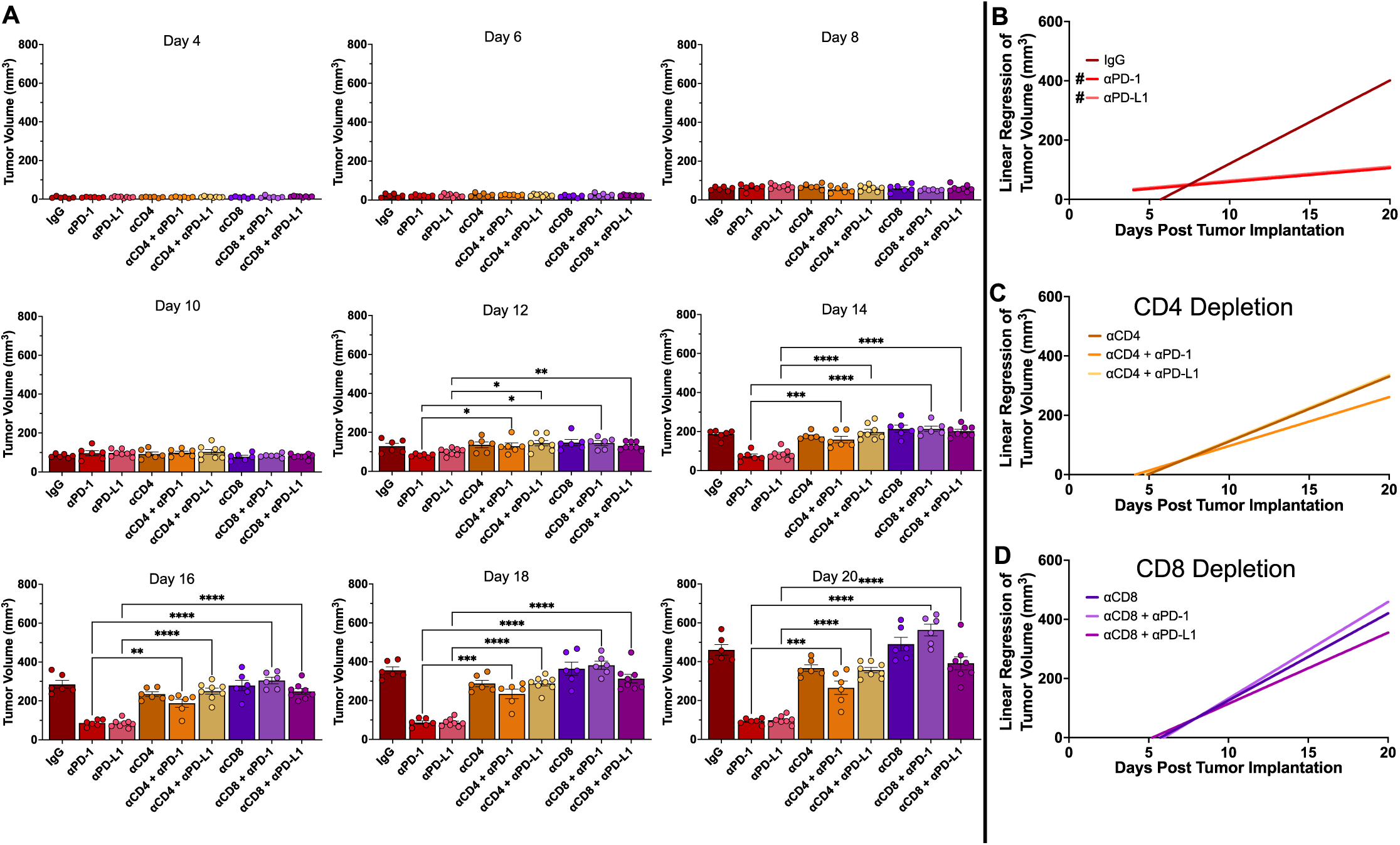
SEMA7A OE tumor growth with **α**PD-1/PD-L1 in the absence of CD4+ or CD8+ T cells. (A) Tumor volumes for SEMA7A overexpressing E0771 tumors in vivo. Respective CD4-and CD8-depleted groups received αCD4 or αCD8 at days 6, 8, 13, and 18. IgG, αPD-1, or αPD-L1 treatments were administered at days 10, 13, 16, and 19. Sample sizes: IgG *n*=6 tumors; αPD-1 *n*=6 tumors; αPD-L1 *n*=8 tumors; αCD4 + IgG *n*=6 tumors; αCD4 + αPD-1 *n*=6 tumors; αCD4 + αPD-L1 *n*=8 tumors; αCD8 + IgG *n*=6 tumors; αCD8 + αPD-1 *n*=6 tumors; αCD8 + αPD-L1 *n*=8 tumors. Error bars shown as mean ± SEM. Statistics are from ordinary one-way ANOVA with Tukey’s multiple comparisons test. *\*p<0.05, **p<0.01, ***p<0.001, ****p<0.0001*. (B-D) Growth rates from Figure 6 A-C.

**Figure S7:**
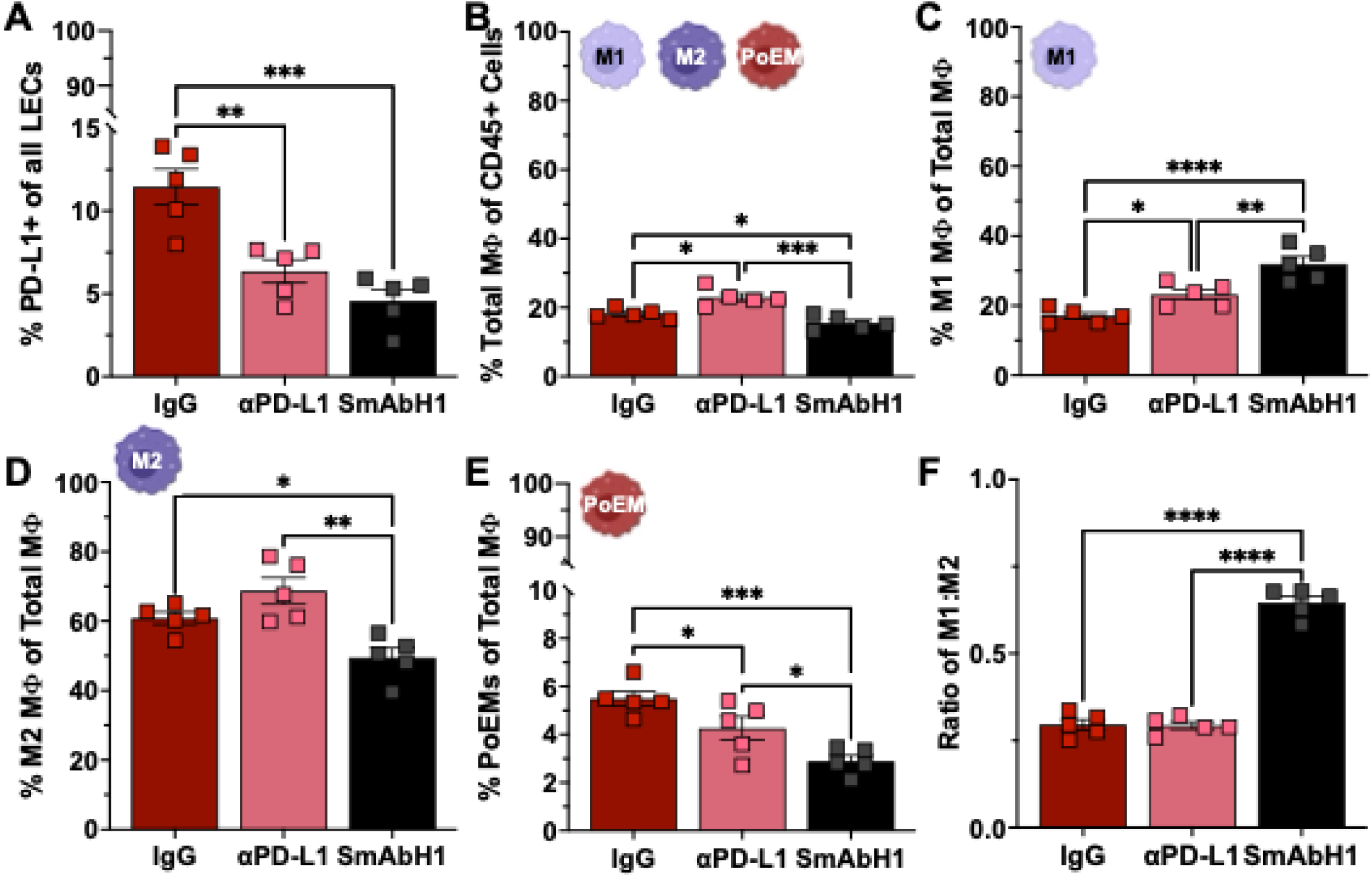
SmAbH1 in vitro binding and killing efficacy and in vivo effects on myeloid cells. (A) ELISA of SmAbH1 binding to purified SEMA7A protein. (B) IncuCyte proliferation analysis of 66cl4 control (Ctrl) and SEMA7A overexpressing (OE) cells in the presence of SmAbH1. Results are from pooled technical triplicates and error bars are mean ± SD and statistics shown are from ordinary two-way ANOVA with Šídák’s multiple comparison test. *\*p<0.05, **p<0.01, ***p<0.001, ****p<0.0001*. (C) Clonogenic growth assay analysis of 66cl4 Ctrl, SEMA7A KD, and SEMA7A OE cells in the presence of SmAbH1. Results are from pooled biological and technical triplicates and error bars are mean ± SD and statistics shown are from ordinary two-way ANOVA with Šídák’s multiple comparison test. *\*p<0.05, **p<0.01, ***p<0.001, ****p<0.0001*. (D) Percent PD-L1+ LECs from tumors following treatments outlined in Figure 7D. (E) Percent total macrophages from tumors following treatments outlined in Figure 7D. (F) Percent M1-like macrophages from tumors following treatments outlined in Figure 7D. (G) Percent M2-like macrophages from tumors following treatments outlined in Figure 7D. (H) Percent PoEMs from tumors following treatments outlined in Figure 7D. (I) Ratio of M1-like:M2-like macrophages from tumors following treatments outlined in Figure 7D. Error bars are mean ± SEM. Sample sizes for C-H: *n*=5 tumors/group. Statistics for C-H are ordinary one-way ANOVA with Tukey’s multiple comparisons test. *\*p<0.05, **p<0.01, ***p<0.001, ****p<0.0001*.

**Figure S8:**
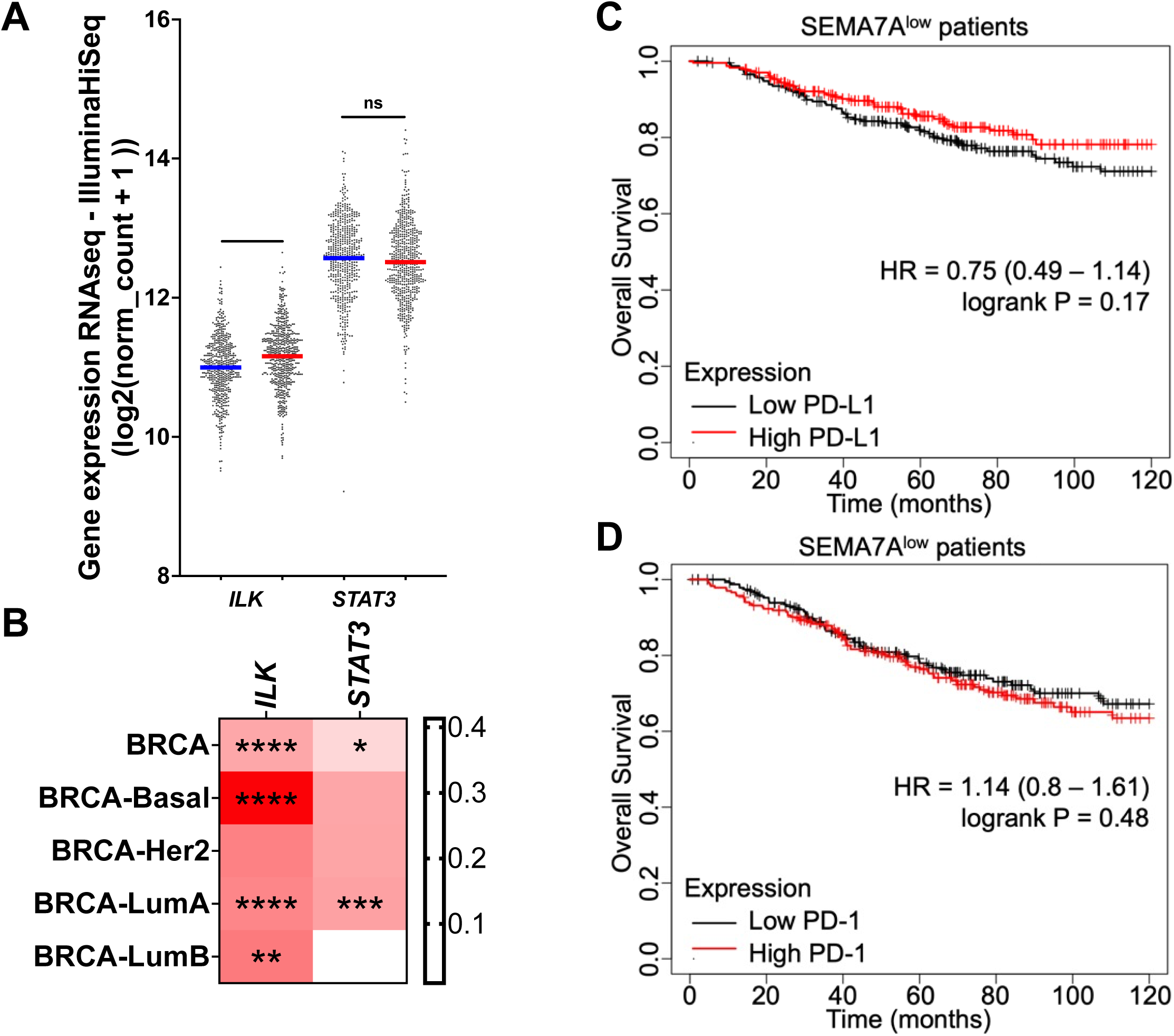
PD-1 and PD-L1 expression do not predict for survival when SEMA7A expression is low. (A) Co-expression analysis from The Cancer Genome Atlas (TCGA) breast cancer dataset using the Xena Functional Genomics Explorer^80^ for *SEMA7A, ILK, and STAT3*. Unpaired two-tailed t test. (B) TIMER analysis^81^ from TCGA breast cancer data set for gene correlations between *SEMA7A* and *ILK or STAT3*. Gene correlations stratified by breast cancer subtypes. Heatmaps generated using purity-adjusted partial spearman’s rank correlation coefficient. (C) Overall survival analysis of SEMA7A^low^ breast cancer patients stratified by high or low expression of PD-L1 using the KM Plotter tool^82^. (D) Overall survival analysis of SEMA7A^low^ breast cancer patients stratified by high or low expression of PD-1 using the KM Plotter tool^82^.

**Figure S9: Flow cytometry gating strategy for mammary epithelial cells, tumor cells, and lymphatic endothelial cells.**

(A) Mammary epithelial cells (normal involution studies in Figure 1) and tumor cells were identified from CD45-/EpCAM+ and lymphatic endothelial cells (LECs) were identified from CD45-/EpCAM-/CD31+/podoplanin+. Cells were further analyzed for SEMA7A or PD-L1 expression.

**Figure S10: Flow cytometry gating strategy for macrophages.**

(A) Macrophages were identified by CD45+/Ly6G-/CD11b+/F4/80+ with M1-like, M2-like, and PoEM macrophages being stratified by MHC II^high^, CD206+, or podoplanin+, respectively, then further characterized by their expression of PD-L1.

**Figure S11: Flow cytometry gating strategy for macrophages.**

(A) T cells were identified from CD31-/EpCAM-/CD45+/B220-/CD3+ and stratified by CD4 or CD8 positivity. CD4+ and CD8+ T cells were then analyzed for activation (PD-1^mid^/Lag-3-/IFNγ+/TNFα+) or exhaustion (PD-1^high^/Lag-3+/IFNγ-/TNFα-).

**Table S1.**
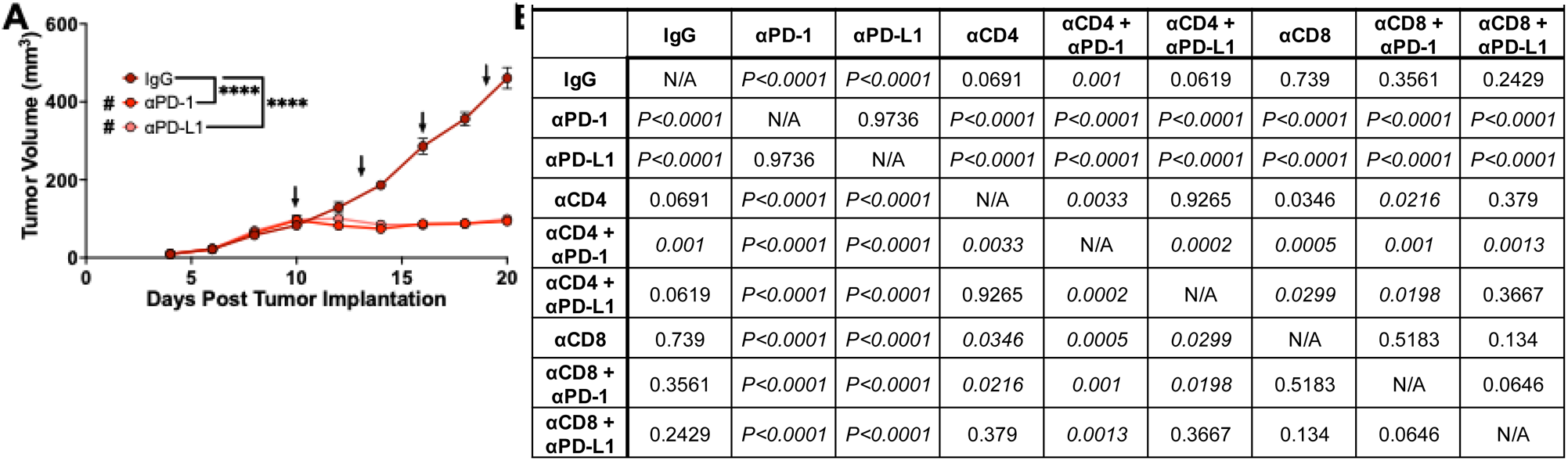
Linear Regression Statistics Between Groups From Figure 5.

| Supplemental Table 1. Linear Regression Statistics Between Groups From Figure 5 |  |  |  |  |  |  |  |  |  |
| --- | --- | --- | --- | --- | --- | --- | --- | --- | --- |
|  | IgG | αPD-1 | αPD-L1 | αCD4 | αCD4 + αPD-1 | αCD4 + αPD-L1 | αCD8 | αCD8 + αPD-1 | αCD8 + αPD-L1 |
| IgG | N/A | $P<0.0001$ | $P<0.0001$ | 0.0691 | 0.001 | 0.0619 | 0.739 | 0.3561 | 0.2429 |
| αPD-1 | $P<0.0001$ | N/A | 0.9736 | $P<0.0001$ | $P<0.0001$ | $P<0.0001$ | $P<0.0001$ | $P<0.0001$ | $P<0.0001$ |
| αPD-L1 | $P<0.0001$ | 0.9736 | N/A | $P<0.0001$ | $P<0.0001$ | $P<0.0001$ | $P<0.0001$ | $P<0.0001$ | $P<0.0001$ |
| αCD4 | 0.0691 | $P<0.0001$ | $P<0.0001$ | N/A | 0.0033 | 0.9265 | 0.0346 | 0.0216 | 0.379 |
| αCD4 + αPD-1 | 0.001 | $P<0.0001$ | $P<0.0001$ | 0.0033 | N/A | 0.0002 | 0.0005 | 0.001 | 0.0013 |
| αCD4 + αPD-L1 | 0.0619 | $P<0.0001$ | $P<0.0001$ | 0.9265 | 0.0002 | N/A | 0.0299 | 0.0198 | 0.3667 |
| αCD8 | 0.739 | $P<0.0001$ | $P<0.0001$ | 0.0346 | 0.0005 | 0.0299 | N/A | 0.5183 | 0.134 |
| αCD8 + αPD-1 | 0.3561 | $P<0.0001$ | $P<0.0001$ | 0.0216 | 0.001 | 0.0198 | 0.5183 | N/A | 0.0646 |
| αCD8 + αPD-L1 | 0.2429 | $P<0.0001$ | $P<0.0001$ | 0.379 | 0.0013 | 0.3667 | 0.134 | 0.0646 | N/A |

## STAR Methods

### RESOURCE AVAILABILITY

#### Lead contact

Further information and requests for resources and reagents should be directed to and will be fulfilled by the lead contact, Traci Lyons, PhD.

#### Materials availability

These studies utilized our novel anti-semaphorin7A monoclonal antibody, “SmAbH1.” There are restrictions to the availability of SmAbH1 due to its pending patent approval and MTA restrictions by the University of Colorado | Anschutz Medical Campus.

#### Data and code availability

- Flow cytometry data reported in this paper will be shared by the lead contact upon request. This paper also analyzes existing publicly available data. These accession numbers for the datasets are listed in the key resources table.
- This paper does not report original code.
- Any additional information required to reanalyze the data reported in this paper is available from the lead contact upon request.

### METHOD DETAILS

#### Cell culture

Authenticated E0771 female mouse mammary carcinoma cells were acquired from CH3 BioSystems and cultured in RPMI 1640 with 10% fetal bovine serum, and 1% penicillin/streptomycin. Authenticated 66cl4-luciferase female mouse mammary carcinoma cells were acquired from Dr. Heide Ford (University of Colorado | Anschutz Medical Campus, Aurora, CO) and cultured in DMEM High glucose with 10% iron-fortified calf serum, 1% MEM non-essential amino acids, and 1% penicillin/streptomycin. 66cl4-luciferase and E0771 cells were transfected with plasmids containing DDK tagged *Sema7a* or a DDK empty vector control plasmids using X-tremeGENE transfection reagent^20^. Transfected cells were selected and maintained using G418 (neomycin and concentration is 0.2mg/ml). *Sema7a* shRNA knockdown and scramble control plasmids were obtained from the University of Colorado Functional Genomics Shared Resource and 66cl4-luciferase cells were transfected in 8ug/ml polybrene and selected for using 3ug/ml puromycin. MDA-MB-231 (obtained from Dr.

Pepper Schedin, Oregon Heath and Sciences University, Portland, OR) and MDA-MB-468 (obtained from Dr. SteveAnderson at the Cell Technologies Shared Resource (CTSR), University of Colorado | Anschutz Medical Campus, Aurora, CO) female breast cancer cells were validated by STR profiling in the CTSR at the University of Colorado | Anschutz Medical Campus, and grown in DMEM High glucose with 10% fetal bovine serum, 1% MEM Amino Acid Solution, and 1 % penicillin/streptomycin. All cells were tested for mycoplasma following thawing from liquid nitrogen using the Lonza MycoAlert Detection Kit, kept at low passage numbers, and allowed to grow for up to 10 passages before new cells were thawed.

#### Animal studies

All animal procedures were approved by the University of Colorado | Anschutz Medical Campus Institutional Animal Care and Use Committee. All mice were housed and bred in a controlled environment. For all studies, *n*=5-6 mice were used for each group and performed in triplicate with representative data shown. Male and female C57BL/6 and BALB/c were obtained from Charles River Laboratories. Male and female *Sema7a^tm1Alk^/*J mice (age 6-8 weeks) (a generous gift from Dr. Alex Kolodkin) were bred to establish our *Sema7a^tm1Alk^/*J colony for involution studies. *Sema7a^tm1Alk^/*J mice were crossed with wild-type C57BL/6 mice to generate *Sema7a* heterozygous mice for involution tumor studies. All mice were genotyped using TransnetYX.

For involution studies, 6–8-week-old female *Sema7a^tm1Alk^/*J and C57BL/6 were bred with BALB/c males. Age-matched nulliparous animals were set aside as controls. Mammary gland involution was initiated in the bred females by force weaning pups at 10 to 14 days post-parturition. For normal involution studies, number 4 mammary glands—excluding the inguinal lymph node—were harvested from nulliparous, involution (Inv) day 2, 3, 4, 6, and 8, and regressed (Reg; involution day 28) mice. For involution tumor studies, 250,000 E0771 cells were injected into the number 4 left and right mammary glands of *Sema7a^tm1Alk^/*J, *Sema7a* heterozygous, or wild-type C57BL/6 mice 1 day after forced weaning (involution groups) or into age-matched nulliparous female mice of each genotype.

For E0771 tumor studies, 250,000 E0771-DDK or E0771-DDK-SEMA7A-overexpressing cells were injected into the number 4 left and right mammary glands of 6–8-week-old female C57BL/6 mice. For 66cl4 tumor studies, 200,000 66cl4-DDK or 66cl4-DDK-SEMA7A-overexpressing cells were injected into number 4 left and right mammary glands of 6–8-week-old female BALB/c mice. Tumors were monitored for signs of ulceration and were measured every other day as soon as they became palpable with tumor volume being calculated using length x width x width x 0.5. E0771 tumors were allowed to grow for 2-3 weeks post implantation for SEMA7A-overexpression, αPD-1/αPD-L1, and T cell-depletion studies, with an exception made for the SmAbH1 survival analysis studies. 66cl4 tumors were allowed to grow for 5-6 weeks post implantation before study end. At the end of all studies, tumors, mammary glands, and lymph nodes were harvested for flow cytometry and immunohistochemistry. For αPD-1/αPD-L1 and SmAbH1 treatment studies, mice were randomized into treatment groups and injected intraperitoneally with 250ug of αPD-1, αPD-L1, SmAbH1, or isotype control (IgG2a, IgG2b, or IgG1, respectively) antibody once tumors became palpable or when average tumor volume reached 100-150mm^3^. For the T cell depletion studies, 200ug of αCD4 or αCD8 antibodies were administered interperitoneally to their respective groups 4 and 2 days (to ensure T cell depletion) prior to the start of αPD-1/αPD-L1 treatment.

#### Tissue dissociation

Number 4 mammary glands (normal involution studies) or implanted tumors (tumor studies) were dissected using micro-scissors taking care to not gather the inguinal lymph nodes. Harvested tissues were placed into 2ml of Click’s media and finely minced with scalpels prior to the addition of 3 ml dissociation buffer consisting of a final concentration of 500units/ml collagenase II (Worthington Biochemical; Cat# LS004174), 500units/ml collagenase IV (Worthington Biochemical; Cat# LS004186), and 20mg/ml DNase I (Worthington Biochemical; Cat# LS002138) per sample. Samples were incubated at 37°C for 1 hour and dissociated tissue suspensions were passed through a 100um filter and washed with 5ml of Isolation Buffer (1x HBSS with 1.6% 30%w/v BSA and 0.4% 500mM EDTA) to gain single cell suspensions. Cell suspensions were then centrifuged at 300g for 5 minutes at 4°C. If the resulting cell pellet possessed significant red blood cells, then cells were incubated with ACK lysis buffer at room temperature for 10 minutes and centrifuged again at 300g for 5 minutes at 4°C. Cell pellets were then resuspended in Isolation Buffer and transferred to 96 well U bottom plates for flow cytometry staining.

#### Flow cytometry

Dissociated mammary glands and tumors were stained for flow cytometric analysis. All flow cytometry panels were stained in 96 well U bottom plates in 100ul total volume consisting of 10ul Brilliant Stain Buffer (Invitrogen; Cat# 00-4409-750), 10ul anti-Fc receptor blocking antibody (2.4G2 anti-CD16/CD32; acquired from Ross Kedl at University of Colorado | Anschutz Medical Campus, Denver, CO), sum of staining antibodies, and brought to 100ul total volume with FACS buffer (1x PBS with 2% FBS). Samples were stained for 30 minutes at 4°C in the dark, centrifuged at 300 g for 5 minutes at 4°C, and washed twice with 100ul FACS buffer. If intracellular staining was necessary, we utilized the BD Cytofix/Cytoperm Fixation/Permeabilization Kit (BD Biosciences; Cat# 554714) according to manufacturer recommendations and intracellular staining was performed with 10ul Brilliant Stain Buffer, 10ul anti-Fc receptor blocking antibody, sum of staining antibodies, and brought to 100ul volume with BD Perm/Wash Buffer. Following all staining and washing, cells were filtered through 30um filters and analyzed on the ZE5-YETI cytometer (UCCC Flow Cytometry Core). All data were analyzed using FlowJo analysis software.

For the epithelial/endothelial/myeloid panel, cells were stained with CD45 (BUV395; BD Biosciences; Cat#564279) (1:300), EpCAM (PE-Cy7; eBioscience; Cat# 25-5791-80) (1:300), SEMA7A (FITC; Abcam; Cat# ab26012) (1:50), PD-L1 (BV711; BioLegend; Cat# 124319) (1:100), CD31 (BUV737; BD Biosciences; Cat# 612802) (1:300), Podoplanin (APC; BioLegend; Cat# 127409) (1:200), Ly6G (BV510; BioLegend; Cat# 127633) (1:100), CD11b (AF594; BioLegend; Cat# 101254) (1:200), CD11c (BV786; BioLegend; Cat# 117336) (1:200), MHC II (Pacific Blue; BioLegend; Cat# 107620) (1:200), F4/80 (PerCP; BioLegend; Cat# 123126) (1:200), and CD206 (BV605; BioLegend; Cat# 141721) (1:200).

For the T cell panel, cells were stained with CD31 (BUV737; BD Biosciences; Cat# 612802) (1:300), EpCAM (BUV737; BD Biosciences; Cat# 741818) (1:300), CD45 (BUV395; BD Biosciences; Cat# 564279) (1:300), B220 (BUV661; BD Biosciences; Cat# 612972) (1:300), CD3 (BV785; BioLegend; Cat# 100232) (1:200), CD4 (APC-Fire810; BioLegend; Cat# 100480) (1:100), CD8a (Pacific Blue; BioLegend; Cat# 100725) (1:200), PD-1 (BV510; BioLegend; Cat# 135241) (1:100), Lag-3 (BV711; BioLegend; Cat# 125243) (1:100), IFNγ (AF594; BioLegend; Cat# 505845) (1:200), TNFα (PE-Cy7; BioLegend; Cat#506323) (1:200).

Mammary epithelial cells and tumor cells were identified from CD45-/EpCAM+ and lymphatic endothelial cells (LECs) were identified from CD45-/EpCAM-/CD31+/podoplanin+ (SFig. 9). Macrophages were identified by CD45+/Ly6G-/CD11b+/F4/80+ with M1-like, M2-like, and PoEM macrophages being stratified by MHC II^high^, CD206+, or podoplanin+, respectively. Tumor cells, LECs, and macrophages were then further characterized by their expression of SEMA7A and/or PD-L1 (SFig. 10). T cells were identified from CD31-/EpCAM-/CD45+/B220-/CD3+ and stratified by CD4 or CD8 positivity. CD4+ and CD8+ T cells were then queried for activation (PD-1^mid^/Lag-3-/IFNγ+/TNFα+) or exhaustion (PD-1^high^/Lag-3+/IFNγ-/TNFα-) (SFig. 11).

#### SEMA7A signaling assays

E0771-DDK, E0771-DDK-SEMA7A, 66cl4-DDK, and 66cl4-DDK-SEMA7A cells were seeded at 4 x 10^4^ per well in 96 well plates and MDA-MB-231 cells were seeded at 3 x 10^5^ cells per well in a 6 well dish and allowed to adhere for 24 hours prior to addition of inhibitors or SEMA7A stimulation. After 24 hours, cells were serum starved for 3 hours followed by preincubation with monoclonal antibodies or inhibitors for 45 minutes as follows; 250nM IgG1 (Bio X Cell; Cat# BE0083), 250nM IgG2a (Bio X Cell; Cat# BE0089), 10ug/ml αintegrin-α6 (eBioscience; Cat# 14-0495-85), 1ug/ml αintegrin-β1 (Human: BD Bioscience; Cat# 552828 | Mouse: BD Bioscience; Cat# 550531), 250nM αSEMA7A SmAbH1 (University of Colorado Cancer Center (UCCC) Cell Technologies Shared Resource), 1uM CPD22 ILK inhibitor (EMD Millipore; Cat# 407331), or 10uM LY294002 PI3K inhibitor (Cell Signaling; Cat# 9901S). Following preincubation with monoclonal antibodies or inhibitors, cells were stimulated with 25ug/ml of purified SEMA7A (UCCC Cell Technologies Shared Resource) for 45 minutes.

Following inhibition and SEMA7A stimulation, E0771 and 66cl4 cells were lifted using 0.05% Trypsin-EDTA, moved to 96 well U bottom plates, and stained for 30 minutes at 4°C with αPD-L1 (BV711: BioLegend; Cat# 124319). Cells were then washed with flow buffer (above) and analyzed on the ZE5-YETI flow cytometer (UCCC Flow Cytometry Core). Experiments were repeated in biologic and technical triplicates. For MDA-MB-231 cells, protein extracts were prepared by lysing treated cells in RIPA with EDTA (Thermo Fisher; Cat# J61529.AP) containing PhosSTOP phosphatase inhibitor (Roche; Cat# 4906837001), and cOmplete protease inhibitor (Roche; Cat# 11697498001). 20ug of total protein in Laemmli protein sampler buffer were heated at 70°C for 10 minutes and analyzed by Western Blot using a 4-20% TGX SDS-gel and transferred to a PVDF membrane. Membranes were blocked with 5% bovine serum albumin and probed with the following primary antibodies according to manufacturer recommendation: pAKT S473 (Cell Signaling; Cat# 4060), total AKT (Cell Signaling; Cat# 4691), pGSK3β S21/S9 (Cell Signaling; Cat# 9331S), total GSK3β (Cell Signaling; Cat# 12456), PD-L1 (Cell Signaling; Cat# 51296 or Abcam; Cat# ab213524), GAPDH (Abcam; Cat# ab9482), and vinculin (Cell Signaling; Cat# 13901). Membranes were washed with 1X Tris-buffered saline with Tween 20, incubated with a goat anti-rabbit HRP secondary antibody (Abcam; Cat# 6721), and developed with enhanced luminescence detection system (Thermo Fisher; Cat# 32106) on the Licor Odyssey. Densitometric O.D. values of three biological replicates were obtained using ImageJ^110^ and normalized to the loading control GAPDH or vinculin.

#### ELISA

96 well MaxiSorp immuno plates (Thermo Fisher; Cat# 436110) were coated with 50ug/ml SEMA7A purified in ELISA Phosphate Coating Buffer (Thermo Fisher; Cat# CB07100) and incubated for 3 hours at room temperature. Plates were then washed once with 1X Wash Buffer (Thermo Fisher; Cat# WB01), blocked with 200ul 1X Assay Buffer (Thermo Fisher; Cat# DS98200), and incubated for 2 hours at room temperature with gentle rocking. After blocking, Assay Buffer was aspirated and fresh 100ul 1X Assay Buffer was added to all wells except the first in the dilution series. Serial dilutions of SmAbH1 were performed by adding 200ul SmAbH1—at a final concentration of 200ug/ml—to well 1 and diluted using a 100ul serial dilution from well 1-9 for final concentrations (ug/ml) of 200, 100, 50, 25, 12.5, 6.25, 3.125, 1.563, and 0.781. Wells 10-12 were used as blank controls. Plates were incubated for 1 hour at room temperature and then overnight at 4°C with gentle rocking. The following morning plates were washed twice with 1X Wash Buffer and 100ul goat anti-mouse IgG HRP 2° (Abcam; Cat# ab97040) in 1X Assay Buffer was added to wells 1-11 with 100ul 1X Assay Buffer alone added to well 12. Plates were incubated for 2 hours at room temperature with gentle rocking and washed 4 times with 1X Wash Buffer prior to developing. After washes, 150ul SuperSignal ELISA Femto (Thermo Fisher; Cat# 37075) was added to each well, incubated for 1 minute with a microplate mixer, and absorbance at 425nm was measured using the BioTek SynergyH1 plate reader and Gen5 software. All samples and controls were plated in duplicates which were used to create standard curve using log transformation X=(Log)X / nonlinear fit in PRISM.

#### IncuCyte proliferation assay

The antiproliferative effect of SmAbH1 was evaluated in 66cl4 SEMA7A DDK cells using the IncuCyte S3 Live-Cell Analysis System (Essen Bioscience). Cells were plated in a 96-well plate in triplicate at 1,500 cells/well and allowed to adhere for 24 hours. After 24 hours, the cells were treated with Mouse IgG1 control (1000nM | Bio X Cell; Cat# BP0083) or SmAbH1 (10nM, 100nM, 200nM, 500nM, or 1000nM). The plate was promptly transferred to the Sartorius S3 IncuCyte to be cultured at 37°C for seven days, and photos were taken every 4 hours. Following seven days, the plate was removed and discarded. Percent confluence was analyzed using the IncuCyte Zoom 2020A software (Essen Bioscience) and normalized to 0hr for each treatment. Experiments were performed in biological and technical triplicates and statistics were measured via repeated-measures two-way ANOVA with multiple comparisons. Statistics shown on the graph were pulled from the last data points from multiple comparisons. Overall ANOVA results p<0.0001.

#### SEMA7A-monoclonal antibody clonogenic assay

66cl4 *Sema7a*-overexpressing, knockdown, or control cells were seeded at 1,000 cells/well in 12-well plates and allowed to adhere overnight at 37°C. The following morning, αSEMA7A SmAbH1 was added for final concentrations of 0nM, 10nM, 50nM, 100nM, 250nM, 500nM, or 1,000nM. Murine IgG1 at a concentration of 1,000nM was used as a control. Cells were incubated for 7 days at 37°C, after which the media was removed, and the cells were washed with sterile PBS. The cells were fixed with 5mL of 10% neutral-buffered formalin solution for 5 minutes. The NBF was discarded, and the plates were stained with 0.1% crystal violet in mqH_2_O for 5 minutes. The stain was discarded, and the plates were washed three times with mqH_2_O and dried overnight. Experiments were performed in biological and technical triplicates. Individual colonies were counted by hand and blinded until counts were done.

#### TCGA RNASeq analysis

We analyzed The Cancer Genome Atlas (TCGA) breast cancer dataset for gene co-expression and correlations of *SEMA7A*, *PDCD1* (PD-1), *CD274* (PD-L1), *PDPN* (podoplanin), *CD68*, and *MRC1* (mannose receptor/CD206). Gene expression analysis was performed using the Xena Functional Genomics Explorer^80^ where patients were stratified by high and low expression of *SEMA7A*—based on the dataset mean expression level—followed by co-expression analysis for the genes of interest. Statistics were measured in the Xena Database using spearman’s rank correlation coefficient. Gene correlation analysis was performed using the TIMER2.0 Gene_Corr module^81^ and heatmaps were generated using the purity-adjusted partial spearman’s rank correlation coefficient.

#### Kaplan–Meier survival analysis

The Kaplan–Meier Plotter is a tool utilized to assess the correlation and effect of all genes on survival using over 35,000 samples from 21 tumor types^82^. Survival statistics were performed using KM Plotter tool on cohorts of tissues. For *SEMA7A* vs *PDCD1/CD274* (PD-1/PD-L1), patients from the mRNA gene chip were stratified by high or low median expression of *SEMA7A* and then compared to high vs low expression of *PDCD1* or *CD274* for overall survival at 10 years. For Figure 8E, the “KM Plotter for immunotherapy” tool was used to query overall survival based on high vs low *SEMA7A* expression across all tumor types (as there were not enough breast cancer patients for analysis) in patients that have received αPD-1 immunotherapy.

#### Quantification and statistical analysis

All quantitative and statistical analyses outside of Xena, TIMER2.0, and KM Plotter were performed using GraphPad Prism and the methods of test are labeled in figure legends. Statistical tests include unpaired t test, one-way ANOVA with Dunn’s multiple comparison test, two-way ANOVA with Šídák’s multiple comparison test, simple linear regression for tumor growth rates, and Kaplan-Meier survival analysis with Mantel-Cox comparison test. Only P values of less than 0.05 were considered significant.

## In brief

Highlights

- We describe a link between SEMA7A signaling and PD-L1 expression.
- SEMA7A+ tumors are enriched for PD-L1 and immunosuppressive cells.
- αPD-1/αPD-L1 treatments enrich for SEMA7A+ tumor cells
- αSEMA7A treatment decreases tumor growth and promotes anti-tumor immunity

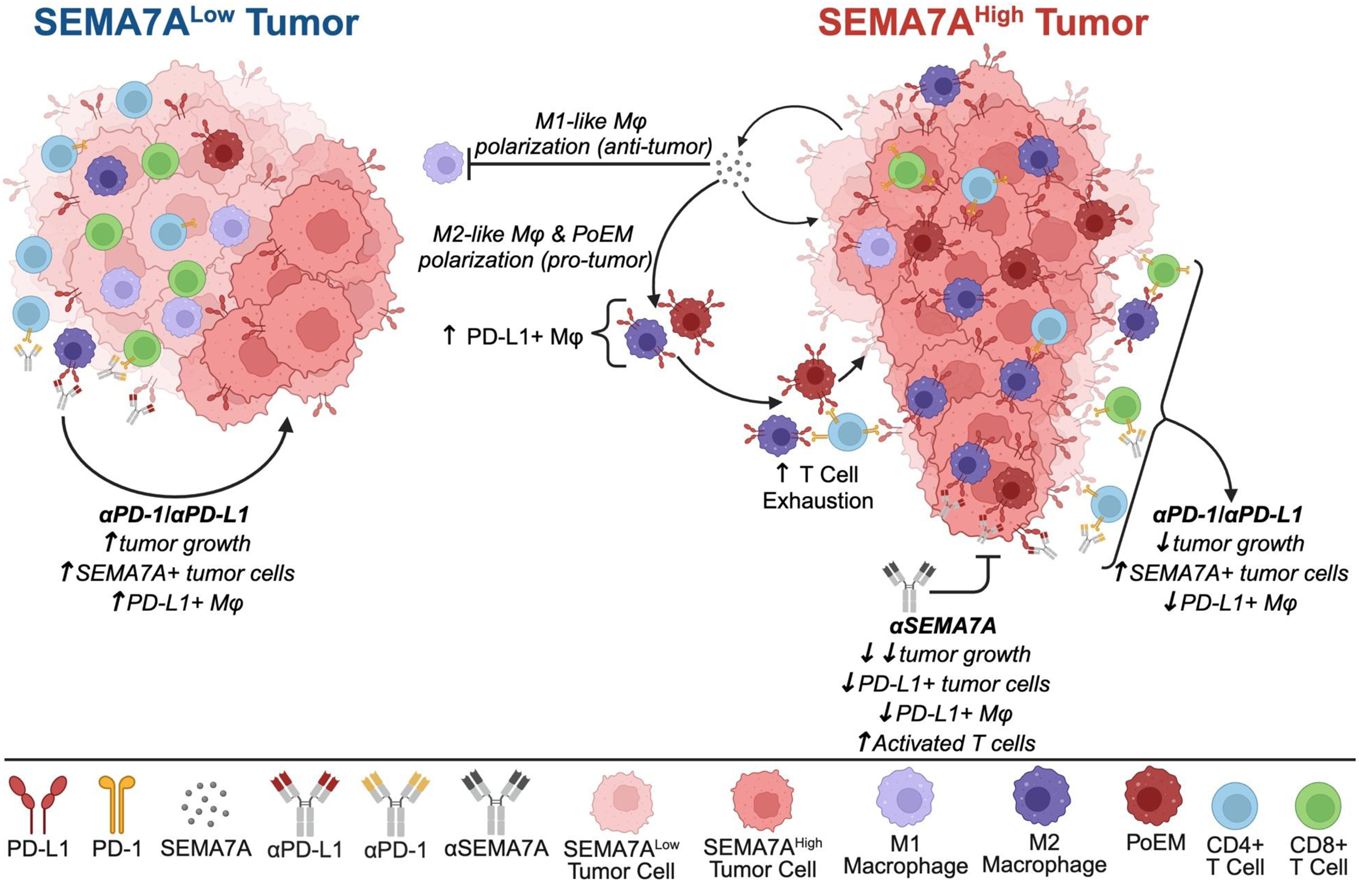

